# Development of a Quantitative Systems Pharmacology Model to Interrogate Mitochondrial Metabolism in Heart Failure

**DOI:** 10.1101/2025.07.08.663697

**Authors:** Lyndsey F. Meyer, Neda Nourabadi, Cynthia J. Musante, Daniel A. Beard, Anna Sher

**Author notes:** Corresponding Author: Lyndsey Meyer. Daniel A. Beard and Anna Sher share senior authorship.

## Abstract

The metabolic hallmarks of heart failure (HF) include diminished ATP hydrolysis potential and alterations in myocardial energy substrate metabolism, such as a switch in substrate utilization away from fatty acid (FA) to carbohydrate oxidation and reduced metabolic flexibility. However, the mechanisms underlying these phenomena and their potential contributions to impaired exercise tolerance are poorly understood.

We developed a comprehensive quantitative systems pharmacology (QSP) model of mitochondrial metabolism to interrogate specific pathways hypothesized to contribute to reductions in reserve cardiac power output in heart failure. The aim of this work was to understand how changes in mitochondrial function and cardiac energetics associated with heart failure may affect exercise capacity. To accomplish this task, we coupled published *in silico* models of oxidative phosphorylation and the tricarboxylic acid cycle with a model of β-oxidation and extended the model to incorporate an updated representation of the enzyme pyruvate dehydrogenase (PDH) to account for the role of PDH in substrate selection.

We tested several hypotheses to determine how metabolic dysfunction, such as a decrease in PDH activity or altered mitochondrial volume, could lead to marked changes in energetic biomarkers, such as myocardial phosphocreatine-ATP ratio (PCr/ATP). The model predicts expected changes in fuel selection and also demonstrates PDH activity is responsible for substrate-dependent switch driven by feedback from NAD, NADH, ATP, ADP, CoASH, Acetyl-CoA and pyruvate in healthy and simulated HF conditions. Through simulations, we also found elevated malonyl-coA may contribute to lower PCr/ATP ratio during exercise conditions as observed in some HF patients.

**Key Points:**

- Exercise intolerance is a hallmark of heart failure in patients with preserved ejection fraction
- We developed a quantitative systems pharmacology modeling approach with the potential to interrogate mitochondrial pathways hypothesized to contribute to exercise intolerance
- The model was developed and evaluated based on simulating in vitro experimental data
- Substrate selection is an emergent property of the model with an increase in ATP demand resulting in a relative increase in the use of carbohydrates to fuel oxidative phosphorylation, an effect driven by feedback regulation of pyruvate dehydrogenase
- The model was used to predict the potential effects of targeted perturbations to key mitochondrial pathways

## 1. Introduction

Heart failure (HF) is a prevalent disease affecting more than 6 million Americans and represents the underlying cause of approximately 1 in 8 deaths due to heart disease in the United States [1]. While many advances have been made in the early detection and treatment of HF, the mortality rate of HF is still 30% within the first 5 years of diagnosis [2]. The etiology of HF can be multifaceted and, for example, can result from pressure overload, ischemia, or diabetic cardiomyopathy. HF with preserved ejection fraction (HFpEF) currently makes up more than half of the heart failure population and potential treatment options are limited to SGLT-2 inhibitors [3].

Despite the multifaceted etiology, there are some key physiological changes that are considered hallmarks of HF. The failing heart is unable to provide sufficient cardiac output to meet the body’s need for oxygen or is only able to do so when the preload filling pressures are pathologically high [4, 5]. In addition, HF patients exhibit exercise intolerance due to pathologically low recruitable cardiac reserves [6-8]. To compensate, the heart may undergo maladaptive cardiac remodeling in which left ventricle hypertrophy and metabolic inflexibility (an increased reliance on carbohydrate oxidation and impaired ability to oxidize fatty acids as a metabolic fuel) can occur [6]. Presently, it is unknown whether metabolic inflexibility is a cause or consequence of HF. Further exploration and understanding of these metabolic changes may lead to advancements in the treatment of HF.

The healthy myocardium oxidizes both fatty acids and glucose to fuel mitochondrial ATP synthesis. In healthy individuals, under basal conditions, fatty acid oxidation contributes between 40 and 60% of total carbon substrate input to the tricarboxylic acid (TCA) cycle to fuel oxidative ATP production, with carbohydrate oxidation contributing roughly ∼20-60% [9, 10]. When faced with a physiological challenge, such as exercise, the healthy heart exhibits a range of metabolic flexibility by an increased substrate preference for glucose oxidation and glycolysis to meet the increased energetic demand [11, 12]. However, in conditions associated with HF with reduced ejection fraction, HFrEF, this metabolic flexibility is compromised [13]. In the myocardium there is an increased reliance on carbohydrates as the predominant fuel source under basal conditions and, thus, does not exhibit the normal range of metabolic flexibility in substrate use [14]. Reduced metabolic flexibility in HF is associated with a derangement of phosphate metabolite (ATP, ADP, and inorganic phosphate (Pi)) levels [15]. One of the clinically accessible biomarkers of the HF progression is the phosphocreatine/ATP (PCr/ATP) ratio. A decline in PCr/ATP ratio has been shown to be associated with HF severity [16, 17] and implicated as a predictor of cardiovascular mortality; specifically, a PCr/ATP ratio less than 1.6 is associated with increased mortality [18]. Additionally, Lopez et al. have shown in a preclinical rodent model [19] that changes in myocardial phosphate metabolites observed in HF may cause systolic mechanical dysfunction.

One strategy to better understand mitochondrial function in HF is to use quantitative systems pharmacology (QSP) modeling. There exists a history of mathematical models describing various pathways of the mitochondria and glycolysis that can be leveraged to characterize mitochondrial substrate selection. An overview of the pathways described by these models is summarized in Fig. 1 [20-32]. One of the early computational models to describe metabolism in an isolated, Langendorff perfused rat heart was developed by Kohn et al. [26] in which the rates of transporters and enzymatic reactions are explored in quantitative detail. Building on the contributions of Kohn and colleagues, *in silico* models describing kinetics and thermodynamically balanced reaction mechanisms for oxidative phosphorylation, the TCA cycle and metabolite transport have been developed by Beard and Wu et al. [21, 25, 33] (data from rat and canine), Cortassa et al. [27, 30, 34] (data from mouse and guinea pig), and Ghosh et al. [35] (data from rat). However, a validated and experimentally constrained *in silico* model describing mitochondria function and consequences of substrate selection has not been explored. In this work to better understand the causes and consequences of mitochondrial dysfunction in HF, we have coupled the model of oxidative phosphorylation and the TCA cycle [25] with a detailed model of β-oxidation from van Eunen et al. [20].

**Figure 1.**
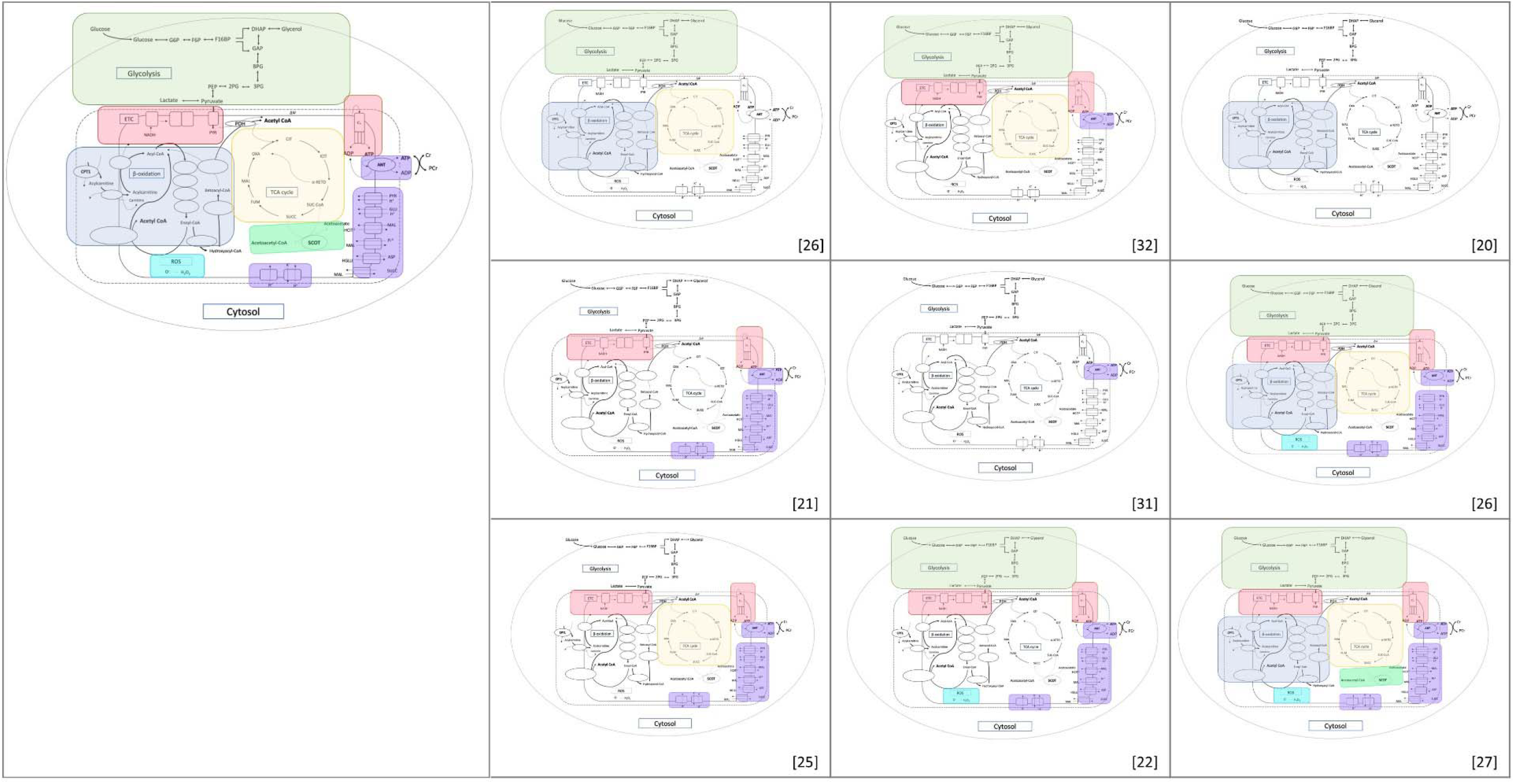
Overview of *In silico* models describing pathways implicated in cardiac energetics. Pathway Key: Glycolysis (green), Electron Transport Chain (Red), TCA Cycle (Yellow), β-Oxidation (light purple), Transporters (Dark Purple, Reactive Oxygen Species (ROS) (Blue), SCOT enzyme (Mint green). *In silico* model scope and summary (top to bottom and left to right): •ODE model with 13 unconstrained parameters describing fatty acid catabolic pathway [26]. •ODE model with 41 parameters and 16 adjustable describing mitochondrial respiration [21]. •ODE model with 31 adjustable parameters describing oxidative phosphorylation and TCA cycle to predict the role of ADP and NADH on TCA flux [25]. •ODE model describing highlighted processes phenomenologically [32]. •ODE model describing the kinetics of ANT transport [31]. •ODE model describing ROS [22]. •ODE model describing competition of β-Oxidation pathway [20] •ODE model phenomenologically describing the fate of lipids in mitochondrial respiration [26]. •ODE unconstrained model phenomenologically describing mitochondrial energetics [27].

First, we demonstrate the newly built, *in silico* model correctly captures data from *in vitro* experiments conducted in isolated rat mitochondria. Then, we incorporate the identified mitochondrial model into an expanded *in silico* model representing mitochondrial dynamics within an intact cell of the myocardium and scaled to represent the whole heart. We explore mechanisms underlying fuel selection using the *in silico* model under resting and exercise conditions. Finally, we investigate pathways in the mitochondria that could potentially explain observed clinical data measuring exercise intolerance in heart failure patients.

## 2. Methods

### 2.1 In Silico Model Development

We built an *in silico* model to investigate ways dysfunction in mitochondrial mechanisms and substrate selection may impact energetics and mechanics in HF. The *in silico* model was developed by integrating the β*-* oxidation model by van Eunen et al. [20], with the TCA cycle and oxidative phosphorylation pathway model from Wu et. al. [33], as illustrated in Fig. 2a. Experimental data from *in vitro* isolated mitochondria, as well as, *in vivo* dog and human data were used to validate *in silico* model predictions. Together these two models represent the two main sources of acetyl-CoA input to the TCA cycle, and thus, provide a framework for simulating the relative contributions of the oxidation of acyl-carnitines versus pyruvate to the production of acetyl-CoA and ultimately ATP availability.

**Figure 2.**
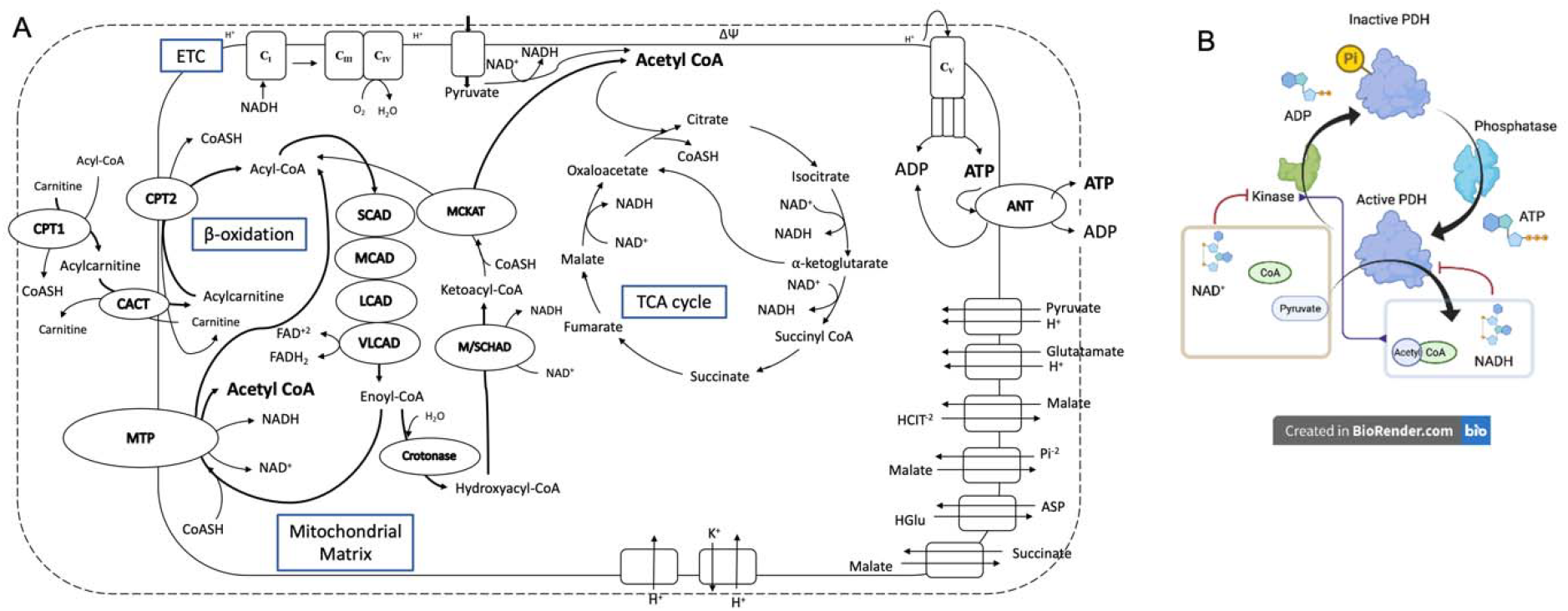
In Silico Mitochondrial Metabolism Model Schematic adapted from Wu et al. [25, 33] and van Eunen et al.[20] **(A)** The mitochondria model characterizes the interplay of the following pathways: (I) Pyruvate enters the mitochondria to generate acetyl-coA (II) β-oxidation converts acyl-carnitines to acetyl-coA and produces NADH and FADH_2_ (III) TCA cycle produces NADH from Acetyl-CoA (IV) electron transport chain (ETC) provides the proton motive force for complex V (C_V_). **(B)** Overview of PDH regulation by pyruvate dehydrogenase kinase (PDK) and pyruvate dehydrogenase phosphatase which are further influenced by the concentrations of acetyl-CoA, NADH, pyruvate, and ATP.

The kinetic operation of the systems represented by these two models are linked via four biochemical reactants: acetyl-CoA (AcCoA), coenzyme A (CoASH), NAD, and NADH. These reactants make up two conserved pools: the total CoA pool and the total pool of NAD and NADH in the mitochondria remain constant over timescales associated with transitions from rest to exercise. AcCoA is a product of *β-*oxidation and pyruvate dehydrogenase (PDH). PDH is a rate limiting enzyme of glucose oxidation and is, therefore, an important mediator of flux through these pathways [36]. NADH, which is the electron donor to complex I of the electron transport chain, is synthesized from NAD by several dehydrogenases in the TCA cycle and *β-*oxidation pathway. NADH also serves as an inhibitor of PDH through activation of its key regulating enzyme, pyruvate dehydrogenase kinase (PDK). Thus, NAD and NADH are important mediators of substrate selection.

To effectively simulate the regulation of relative consumption of fatty acid versus pyruvate, the Wu et al. *in silico* model was further modified to incorporate the regulation of PDH via inactivation by PDK and activation by pyruvate dehydrogenase phosphatase. PDK, in turn, is upregulated by the products of PDH, Acetyl-CoA, and NADH, as well as ATP and downregulated in the presence of excess NAD, CoASH, and pyruvate [37]. The modified structure of regulation of PDH is illustrated in Fig. 2b where the ratios of ATP/ADP, Acetyl-CoA/CoASH, NADH/NAD and pyruvate directly influence the activity of PDH. PDH is also under direct allosteric regulation by AcCoA and NADH. To assess the contributions of PDH and *β -*oxidation to ATP production, the percent of carbohydrate oxidation is quantified by comparing the ratio of flux through PDH (J_PDH_) to flux through citrate synthase (J_CS_). A J_PDH_/J_CS_ ratio of 1 is equivalent to 100% of oxidized substrate contribution by the PDH pathway, i.e. pyruvate. The values of Vmax for both PDH and citrate synthase were adjusted to facilitate model predictions that capture the observed range of values for J_PDH_/J_CS_ ratio reported in the literature [38, 39].

The complete MATLAB (v.2019b, Mathworks, Natick, Ma) model source code is available at https://github.com/pfizer-opensource/mitochondria-metabolism [40]. Additional details about the newly developed *in silico* model can be found in the Appendix.

### 2.2 Respirometry Experiments

#### Isolated Mitochondria Preparation

All the protocols involving animals conformed to the National Institutes of Health Guide for the Care and Use of Laboratory Animals and were approved by the University of Michigan Animal Research Committee. Briefly, adult male Sprague Dawley rats ranging from 300-600 g were anesthetized with an intraperitoneal injection of 0.3–0.5 mL ketamine (90 mg/kg) and 0.3–0.5 mL dexmedetomidine (0.5 mg/kg) followed by 0.5 mL heparin (1000 USP units/mL). Following the protocol of Vinnakota et al., left ventricular tissue was excised and homogenized using a glass dounce homogenizer at 4°C for 3 minutes in isolation buffer (200 mM Mannitol [M9647; Sigma], 60 mM sucrose [S7903; Sigma], 5 mM KH2PO4 [P5379; Sigma], 5 mM MOPS [M1254; Sigma], 1 mM EGTA [E4378; Sigma], and 3 mg of Nagarse protease [P8038; Sigma]). Isolation buffer with 0.1% w/w BSA (A6003; Sigma) was added to the tissue homogenate. The homogenate was then centrifuged at 8000 × g for 10 min at 4°C. Next, the pellet was resuspended in isolation buffer with BSA and centrifuged at 8000 × g for 10 min at 4°C. The resulting pellet was resuspended in isolation buffer with BSA and centrifuged at 700 × g for 10 min at 4°C. The supernatant was collected and centrifuged at 8000 × g for 10 min. Suspensions of purified mitochondria were prepared from ventricular myocardium of male Sprague Dawley rats, following the procedures detailed in Vinnakota et al. [41].

#### Mitochondrial Respiration Measurements

Mitochondrial respiration was measured from purified mitochondria using a high resolution Oroboros Oxygraph 2K, Oroboros (Instruments Gmbh, Innsbruck, Austria). Mitochondria are added to the instrument chamber at a concentration of 0.337 U of citrate synthase activity per mL of respirometer chamber volume, corresponding to approximately 0.1 mg of mitochondrial protein per chamber mL. The respiration buffer contained 5 mM inorganic phosphate (KH2PO4), 90 mM KCl, 1 mM EDTA, 50 mM MOPS, and 0.1% BSA at pH 7.4. All experiments were conducted at 37° C. Mitochondria were incubated for three minutes in respiration buffer prior to adding the substrates. To initiate leak state (state 2) respiration, 20 μM of acyl-carnitine and 2 μM of malate were added simultaneously. After three minutes of leak-state respiration data were recorded, 0.5 mM final concentration of ADP was added to initiate the maximum rate of oxygen consumption in state-3 oxidative phosphorylation (OXPHOS). The maximum rate of steady-state-3 oxygen consumption was recorded for respiration using the following acyl-carnitines: C2: ALC (Acetyl-L-Carnitine), C4: BLC (Butyryl-L-Carnitine), C6: HLC (Hexanoyl-L-Carnitine), C8: OLC (Octanoyl-L-Carnitine), C10: DLC (Decanoyl-L-Carnitine), C12: LLC (Lauroyl-L-Carnitine), C14: MLC (Myristoyl-L-Carnitine), and C16: PLC (Palmitoyl-L-Carnitine).

### 2.3. In Silico Model Calibration and Validation

The β-oxidation model of van Eunen et al. [20] is calibrated by fitting data obtained from experiments on isolated rat liver mitochondria. To adapt the model to represent cardiac mitochondria, model simulated oxygen consumption was compared to the rat data from the respirometry experiments using different chain length acyl-carnitine substrates. Adjusting the *β-*oxidation enzyme Vmax parameters achieved the best fit to the data, yielding the parameter value estimates listed in Table 1.

**Table 1.**
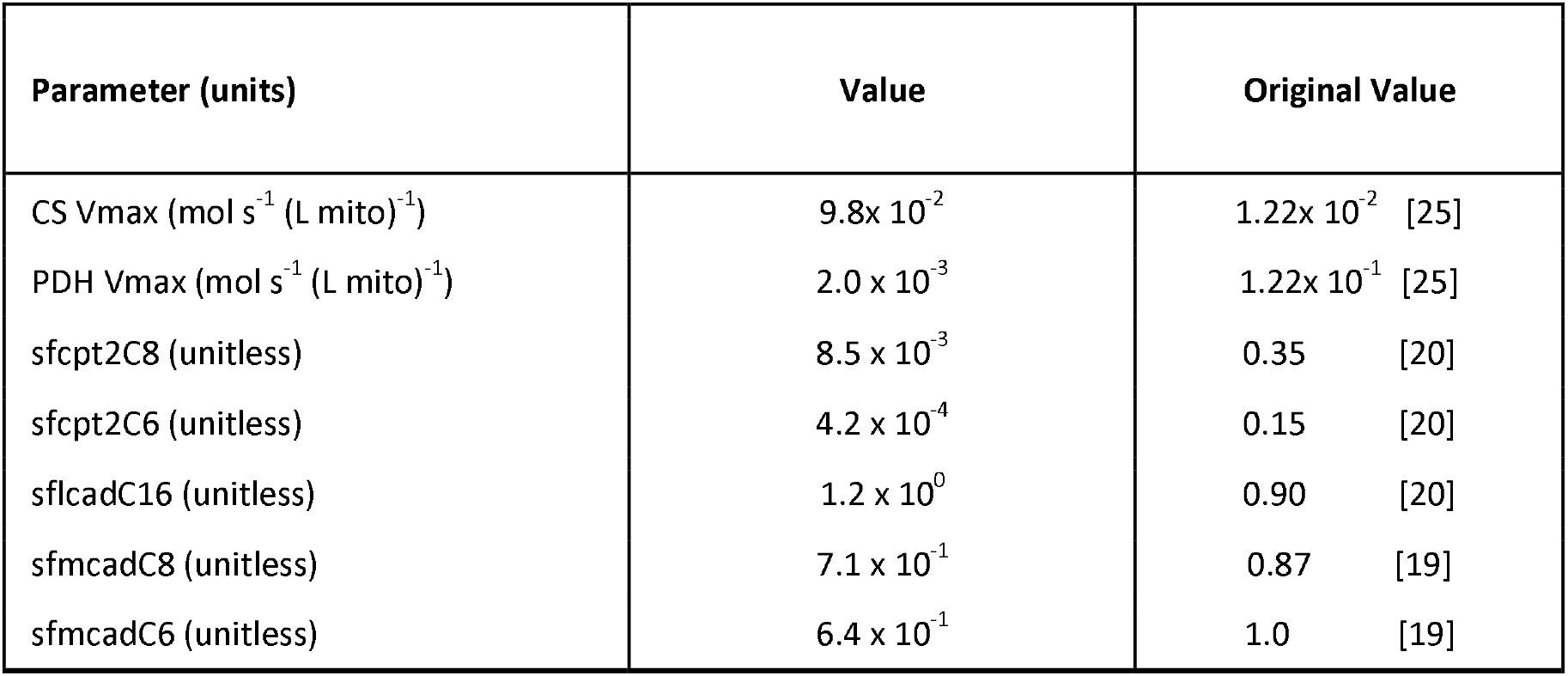
Fitted QSP model parameter values. Remaining parameter values are identical to values provided by van Eunen et al. [20] and Wu et al. [25].

Next, we performed simulations of previously published experimental data found in literature [39] to confirm that the newly developed *in silico* model could recapitulate data beyond the experimental data used to develop the model [38, 42, 43]. Experimental mitochondria respiration data from isolated, mouse mitochondria from cardiac and skeletal muscle [42] were digitized using Web Plot Digitizer version 4.4 (WebPlotDigitizer, 2020, https://automeris.io/WebPlotDigitizer). Additional details of the experimental methods are outlined in [42]. The *in silico* model was initialized to represent the *in vitro* experimental setup, including the addition of extra-mitochondrial substrates malate, glutamate, phosphocreatine and the model predicted oxygen consumption rate, ADP, and PCr/ATP ratio were compared to the digitized data.

### 2.4 Substrate Selection

The same *in silico* mitochondrial model is adapted to represent mitochondrial metabolism of an intact heart by expanding the isolated mitochondrial model to a model of ventricular cardiomyocyte energetics. The representation of ventricular cardiomyocytes includes descriptions of creatine kinase, adenylate kinase, ATP hydrolysis activity in the extramitochondrial space. Fixed boundary conditions exist such that substrates such as pyruvate and palmitoyl-coA are simulated at fixed concentrations which is altered depending on the fasted or fed state. Respirometry experiments use a dilute suspension of isolated mitochondria, therefore, in the *in silico* representation of *in vitro* experiments the external buffer volume is ∼1000 times greater than the mitochondrial volume. For simulations of intact myocardium, it is assumed that mitochondria occupy roughly 28% of cardiomyocyte volume, as in the model of Wu et al. [44]. Finally, for all conditions reported, the *in silico* model for intact myocardium is simulated until steady state is achieved.

To investigate the behavior of the model under varying substrate availability, we simulated the *in silico* model to steady-state with concentrations from 1.2 x 10^−5^ to 1.2 x 10^−2^ M pyruvate and 2.68 x 10^−6^ to 2.68 x10^−3^ M palmitoyl-CoA (ranging from 0.1-fold to 100-fold of the baseline value) under resting, physiological conditions. The resulting flux through PDH (J_PDH_) and CS (J_CS_) was calculated from the model simulations and the ratio was plotted against the changing concentrations for each substrate. Then we simulated the steady-state values of PCr/ATP, and J_PDH_/J_CS_ and Pi accumulation ranging from the resting state to vigorous exercise. Exercise is replicated by varying the myocardial ATP hydrolysis rate (additional details on ATPase flux (J_ATPase_) can be found in the Appendix) from 0.5 mM/s at rest to 1.5 mM/s where 1.5 mM/s represents vigorous exercise [45]. The impact of increasing ATP hydrolysis on substrate selection and energetics is plotted as a function of the free energy of ATP hydrolysis (ΔG_ATP_).

### 2.5 In Silico Model Parameter Uncertainty Assessment

van Eunen et al. [20] and Wu et al. [25] performed local sensitivity analyses on the parameters of the computational β-oxidation and TCA/oxidative phosphorylation models, respectively. The authors report the ranked sensitivity of each parameter. In this work, to explore the uncertainty of *in silico* model projections for PCr/ATP ratio, substrate selection and phosphate accumulation, we generated a parameter set to vary that consisted of the previously identified topmost sensitive parameters from each model. Specifically, for our analysis we chose to include 7 parameters: the activity parameters of PDH, malate dehydrogenase, adenine nucleotide transporter (ANT), and proton leak, the inhibition constant of NADH for α-ketoglutarate dehydrogenase from the TCA/oxidative phosphorylation model, as well as CPT1 transporter specificity and Km from the β-oxidation model. Each parameter was varied by ±20% of the optimized parameter value. To ensure simulations adequately capture the range of parameter space, 100 parameter sets were generated where values for each new parameter set are selected using a Latin hypercube sampling method. Figs. 4B-D illustrate the variability observed in energetic measures across this parameter range where the shaded regions represent the 90th percentile prediction interval.

#### Interrogating Mitochondrial Dysfunction Pathways

Next, we evaluate examples of energetic deficiencies resulting from mitochondrial dysfunction identified from the literature to be implicated in HF. These pathways include the activity of PDH, CPT-1, functioning mitochondrial density, and available pools of adenine nucleotides. Mitochondrial dysfunction pathways are represented in the following cases:

##### Case 1

The activity of PDH and β-oxidation is downregulated by reducing the Vmax of PDH and β-oxidation enzymes by 85% [46] and 90%, respectively.

##### Case 2

The density of mitochondria per unit cell volume is decreased compared to the healthy normal volume, 0.056 L mitochondria/ L cell volume; specifically, the mitochondrial volume is reduced by 20% to simulate an impaired capacity to synthesize ATP in heart failure conditions[47].

##### Case 3

The metabolite pools of total adenine nucleotide and creatine are altered compared to the healthy case.The diminished pools are similar to levels associated with HF and aging [44, 47].

For all cases and conditions, we compared the model-predicted intact myocardium PCr/ATP ratio, J_PDH_/J_CS_ ratio and Pi concentration to the healthy control case. In each simulated case of heart failure, we also explored the impact of interventions proposed to mitigate the loss of energetic capacity, both at rest and under exercise conditions. The following interventions were investigated implicitly in the model through the following modifications:

○ DCA treatment is captured in the model by clamping the PDH in the active (dephosphorylated) state.
○ Nicotinamide riboside supplementationis simulated by elevating NAD levels to increase efficiency of electron transfer to the electron transport chain.
○ Treatment with trimetazidine, an in inhibitor of medium-chain 3-ketoacyl-CoA thiolase (mckat), is simulated by fixing the Vmax of mckat enzyme to 0.

### 2.6 Prediction of Exercise Capacity

Exercise intolerance is a key hallmark of HF. One of the features of exercise intolerance is a decrease in VO_2_ max or the maximum mass of oxygen consumed per unit time, per kilogram of body weight. The VO_2_ max measurement is not a surrogate for the mitochondrial dysfunction since it incorporates whole-body O_2_ related pathways (e.g., Hb levels, O_2_ delivery due to blood flow, etc.), though, impaired mitochondrial function is a contributor to reduced VO_2_ max in HF [48]. In HFrEF, for example, reserve peripheral oxygen consumption is reduced primarily due to impairments in cardiac mechanical function, whereas, in heart failure with preserved ejection fraction it is postulated that impaired oxygen transport and/or consumption in peripheral muscle is a contributing cause of reduced VO2 max [49, 50].

In simulating the effects of myocardial metabolic perturbations on exercise capacity, our underlying assumption is that impaired ATP production by cardiac muscle may reduce cardiac power reserve and would thereby result in measurable reduction in whole-body VO2 max. Thus, in each case of mitochondrial dysfunction, we explored the extent to which maximal myocardial oxygen consumption rate was decreased compared to the healthy case.

### 2.7 Statistics

For results reported in Fig. 5, statistical significance is designated using students t-test with p value < 0.05.

## 3. Results

### 3.1 Model Calibration and Validation

Results of fitting the *in silico* model to measured steady-state oxygen consumptions rates in suspensions of purified mitochondria respiring on different acyl-carnitine substrates are shown in Fig. 3A-B. The observed experimental data are represented by the black bars and the model outputs represented by gray bars. In accordance with the observed data, the *in vitro* model-simulated VO_2_ with increasing carbon chain length of acyl-carnitines. This trend is present for experiments both with and without L-carnitine in the media. Quantitatively, the model predictions fall within one SD of the observed data suggesting the model is in good agreement with the observed data and thus the model of β-oxidation is representative of function in myocardial mitochondria.

**Figure 3.**
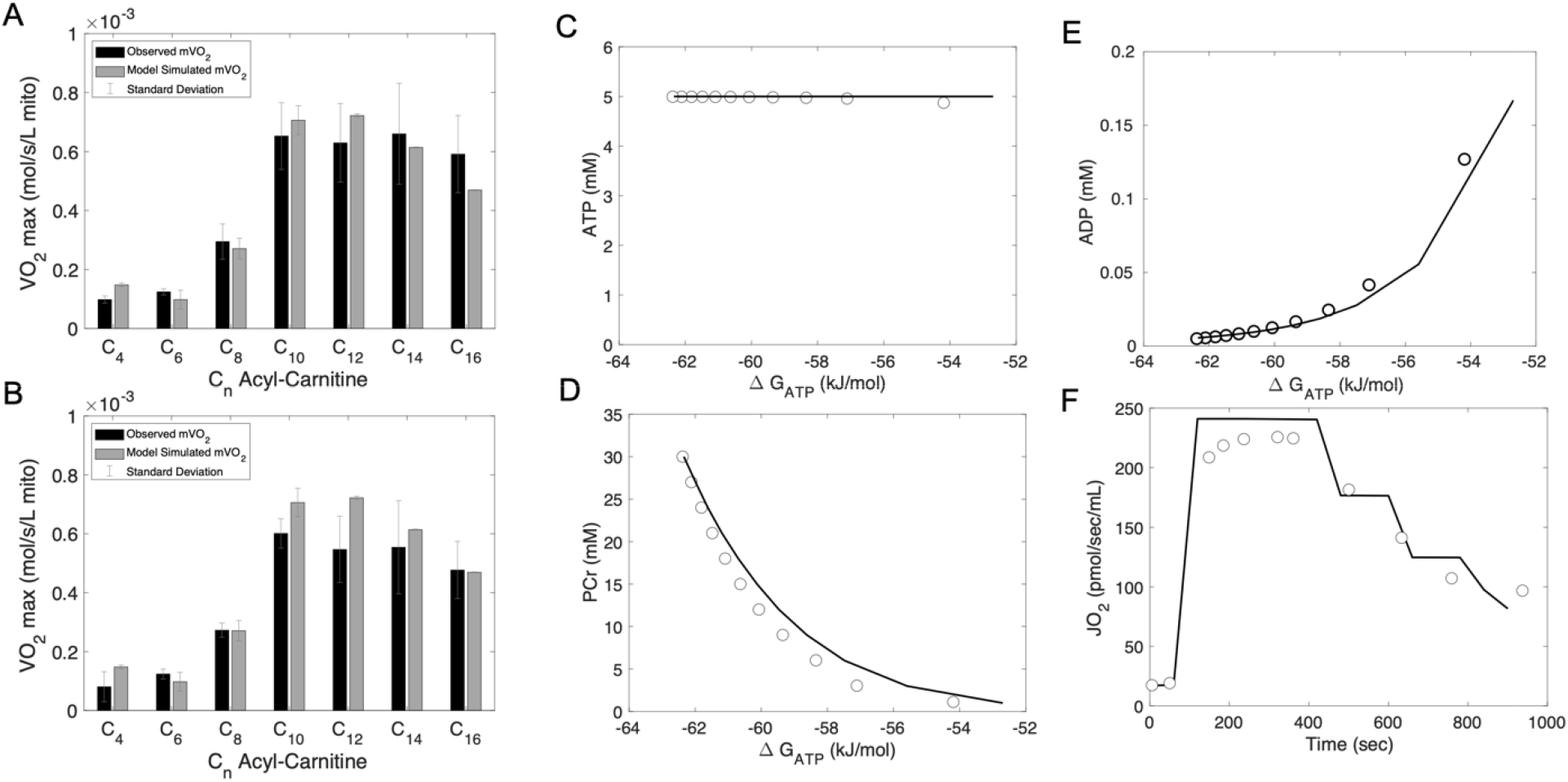
Data Fitting and Model Validation. **(A)** Observed mitochondrial oxygen consumption (mVO_2_) (black bars) supplied with C_n_ Acyl-carnitine in media without L-carnitine present compared with *in silico* model fitted mVO_2_ (gray bars). **(B)** Observed mitochondrial oxygen consumption (mVO_2_) (black bars) supplied with C_n_ Acyl-carnitine in media including the addition of L-carnitine compared with *in silico* model fitted mVO_2_ (gray bars) **(C)** Fixed observed ATP concentrations over time (open circles ) and model simulation (solid line) **(D)** Added phosphocreatine (PCr) concentrations over time (open circles ) and model simulation of PCr addition (solid line) **(E)** Observed change in ADP concentration (open circles) and model predicted change in ADP (solid line) **(F)** Observed change in mitochondrial oxygen consumption (mVO_2_) (open circles) and model predicted mitochondrial oxygen consumption (solid line) following stepwise additions of PCr.

The *in silico* mitochondrial model was validated in part based on comparison to *in vitro* experimental data from isolated mouse mitochondria [42] which were digitized using Web Plot Digitizer (https://apps.automeris.io/wpd/) and simulated as described in the Methods section. The model simulations corresponding to this CK clamp experiment correctly capture steady-state trends of ADP and oxygen consumption following the sequential addition of PCr (Fig. 3). Fig. 3C shows ADP concentrations increasing with increasing ΔG_ATP_ as expected for increasing ATP demand. Similarly, the *in vitro* model quantitatively predicts the observed trend of reducing oxygen consumption ratewith each sequential addition of PCr (Fig. 3D).

### 3.2 Substrate Selection and Energetics in Healthy Cardiomyocytes

To investigate the consequence of substrate availability on substrate selection, we ran the *in silico* model representing intact myocardium and simulated a range of initial concentrations of available palmitoyl-CoA and pyruvate in the extra-mitochondrial cytosolic compartment and observed changes in the predicted J_PDH_/J_CS_ ratio. Fig. 4A demonstrates that in fasted conditions (where concentrations of pyruvate are low, i.e. < 4.0 mM and concentrations of palmitoyl-CoA are high, i.e. > 0.26 mM) there is a strong preference to oxidize fatty acids. In the fed state (high availability of pyruvate and low concentrations of palmitoyl-CoA) preference shifted toward carbohydrate oxidation indicated by the higher J_PDH_/J_CS_ ratio (Fig. 4A). Under these conditions, the PCr/ATP ratio did not change with changing substrate availability (not shown). In addition to investigating a range of substrate concentrations, we simulated substrate selection, PCr/ATP ratio and Pi accumulation from the resting state to vigorous exercise. Vigorous exercise is represented in the model by increasing the rate of ATP hydrolysis from 0.3 mM/s to 1.3 mM/s which corresponds to calculated ΔG_ATP_ ranging from -67 to -59 kJ/mol [44, 51]. Fig. 4B shows that the preferred substrate in the exercise condition is pyruvate when available. Additionally, the PCr/ATP ratio declines with vigorous exercise (Fig. 4C).

**Figure 4.**
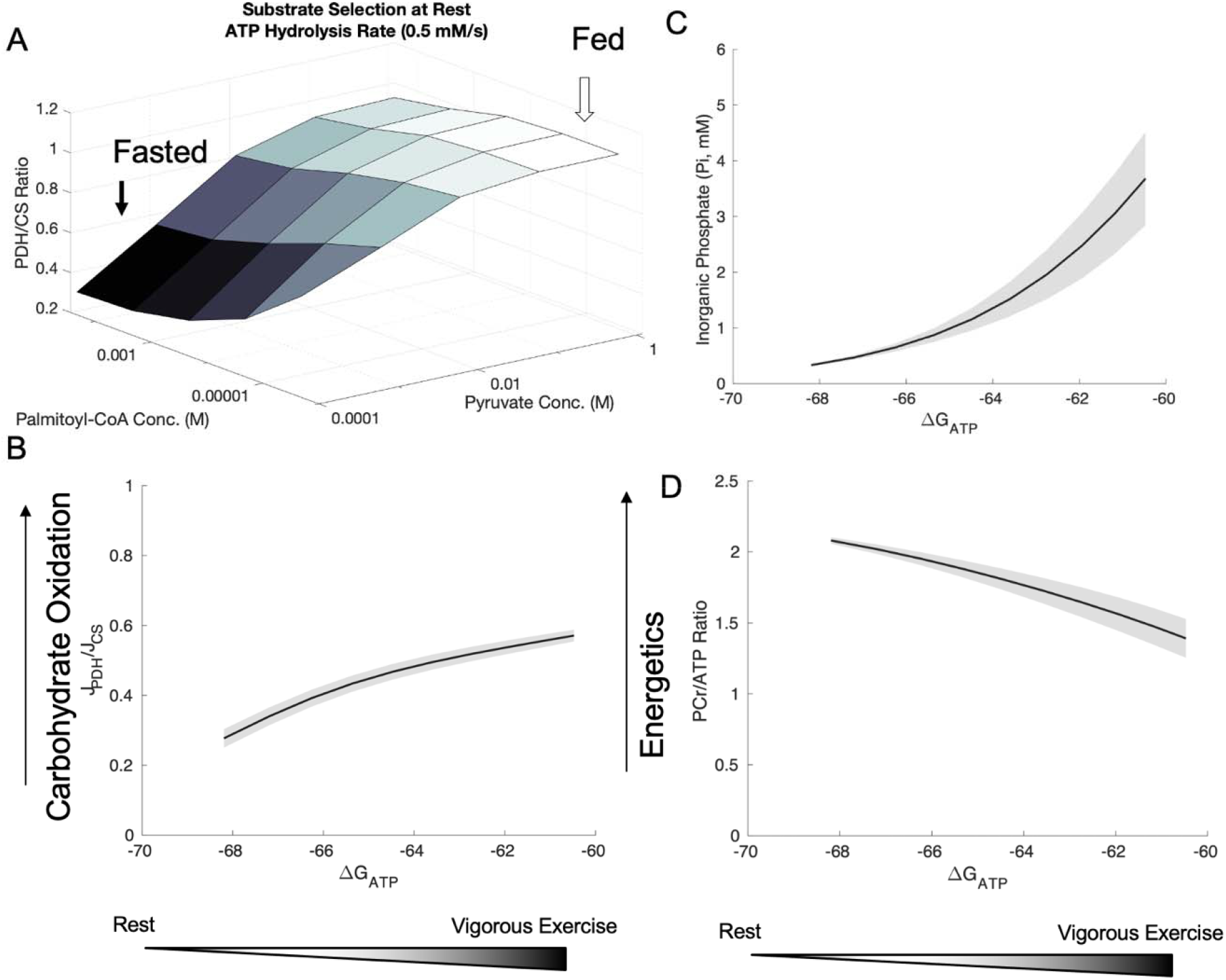
Substrate selection and energetics in healthy cardiomyocytes. Solid line represents the mean values and the shaded region represents the 90% prediction interval. **(A)** Carbohydrate oxidation predominates with increasing pyruvate concentrations illustrating the switch in substrate selection under the fed condition (light gray). In the fasted condition (dark gray) fatty acid metabolism predominates to supply ATP production. **(B)** Fraction of carbohydrate oxidation trends upward with increased free energy of ATP hydrolysis (ΔG_ATP_) from rest to exercise. **(C)** Inorganic phosphate (pi) also trends upward with increased free energy of ATP hydrolysis (ΔG_ATP_) from rest to exercise. **(D)** PCr/ATP ratio declines from 2.1 to 1.5 with increased free energy of ATP hydrolysis (ΔG_ATP_) from rest to exercise.

### 3.3 Substrate Selection and Energetics in HF States

*In silico* results corresponding to simulations of HF states and interventions on intact myocardium are shown in Fig. 5 and summarized in Table 2.

**Figure 5.**
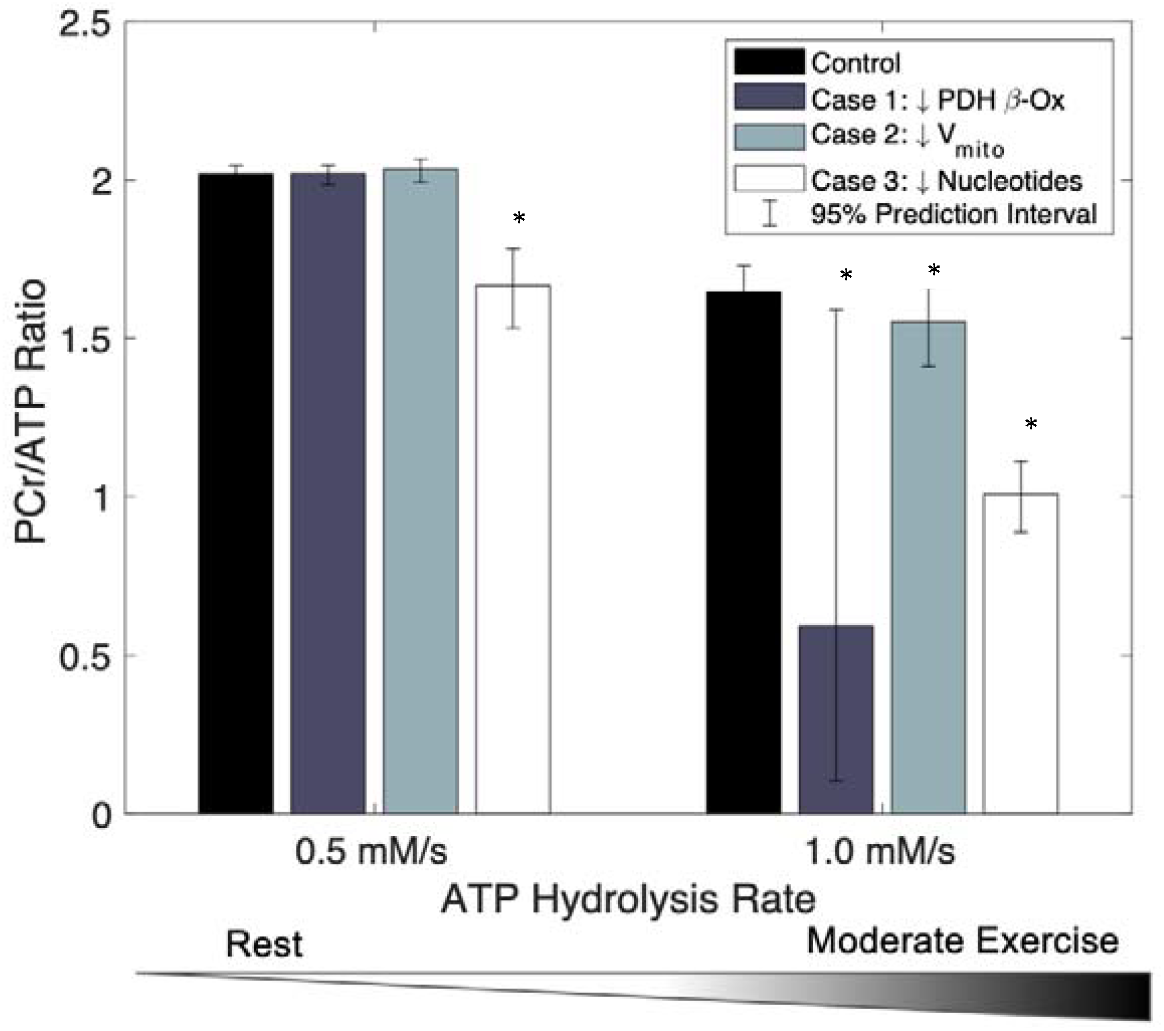
Contribution of pathways to PCr/ATP ratio within each case study. Error bars represent the 95% prediction intervals. Fractional PCr/ATP ratio is shown compared to the healthy control (black bar), reduced adenine nucleotide pools (AMP, ADP, ATP) (dark gray bar) downregulated β-oxidation and PDH pathways (light gray bar), reduced functional mitochondrial content (white bar). Fractional changes in PCr/ATP for each pathway are more pronounced under the exercise conditions compared to rest conditions. * p < 0.05, t-test with respect to control for each group.

**Table 2.** Energetic measures in different cases of HF. Predicted energetic measures for healthy cardiomyocytes and three proposed scenarios representing HF at rest and exercise. Case 1 represents a decrease in PDH activity and β-oxidation enzyme activity. Case 2 represents a 20% decrease in functional mitochondria volume. Case 3 represents a decline in available adenine nucleotide pools.

| Cause of Mitochondrial Dysfunction | Healthy |  | HF Case 1 |  | HF Case 2 |  | HF Case 3 |  |
| --- | --- | --- | --- | --- | --- | --- | --- | --- |
| PDH Activity | - |  | ↓ 85% Vmax |  | - |  | - |  |
| β-Oxidation | - |  | ↓ 90% Vmax |  | - |  | - |  |
| Mitochondria Density | - |  | - |  | ↓ 20% Mito volume |  | - |  |
| Pi, PCr, ATP, NADH pools | - |  | - |  | - |  | ↓ |  |
| Interventions | -ΔG<sub>ATP</sub><br>PCr/ATP<br>MVO<sub>2</sub><br>(μmol/min/g)<br>Rest \| Exercise |  | -ΔG<sub>ATP</sub><br>PCr/ATP<br>MVO<sub>2</sub><br>(μmol/min/g)<br>Rest \| Exercise |  | -ΔG<sub>ATP</sub><br>PCr/ATP<br>MVO<sub>2</sub><br>(μmol/min/g)<br>Rest \| Exercise |  | -ΔG<sub>ATP</sub><br>PCr/ATP<br>MVO<sub>2</sub><br>(μmol/min/g)<br>Rest \| Exercise |  |
| No Intervention (Baseline) | 64.6<br>2.20<br>4.68 | 60.5<br>1.72<br>9.03 | 64.7<br>2.20<br>4.67 | 57.3<br>1.19<br>8.54 | 64.1<br>2.16<br>4.58 | 59.8<br>1.63<br>8.93 | 64.6<br>1.53<br>4.20 | 61.0<br>1.19<br>8.13 |
| Dichloroacetate (DCA)<br>(Maximal flux through PDH<br>and no feedback from PDK) | 66.0<br>2.26<br>4.67 | 62.0<br>1.91<br>9.04 | 67.0<br>2.30<br>4.68 | 62.8<br>1.99<br>9.05 | 65.5<br>2.24<br>4.63 | 61.3<br>1.83<br>9.00 | 65.8<br>1.60<br>4.67 | 62.4<br>1.32<br>9.04 |
| Nicotinamide Riboside (NR)<br>(Increased total NAD<sup>+</sup> pools<br>by 50%) | 64.6<br>2.18<br>4.67 | 60.5<br>1.73<br>9.03 | 64.6<br>2.19<br>4.65 | 57.5<br>1.23<br>8.68 | 64.1<br>2.16<br>4.62 | 59.8<br>1.64<br>8.98 | 64.6<br>1.53<br>4.66 | 61.1<br>1.20<br>9.03 |
| Trimetazidine (Tri)<br>(Inhibitor of mckat,<br>fixed Vmax to 0) | 64.0<br>2.14<br>4.64 | 59.7<br>1.60<br>8.95 | 64.2<br>1.11<br>3.62 | 46.5<br>0.01<br>3.49 | 63.5<br>2.11<br>4.59 | 59.0<br>1.50<br>8.89 | 64.0<br>1.50<br>4.64 | 60.2<br>1.10<br>8.98 |

#### Case 1

Mitochondrial dysfunction in heart failure can significantly increase the degree of reliance on carbohydrate oxidation compared to the healthy case. In this instance, both PDH and β-oxidation enzyme activities are downregulated. Consequently, PCr/ATP declines below a physiological level (ratio of 1.0) at exercise associated with a 1.0 mM/s ATP hydrolysis rate (Fig. 5). Simulating DCA treatment by restoring the PDH activity re-establishes the PCr/ATP ratio to normal levels (Table 2). Under resting conditions, there is no change in the PCr/ATP ratio. Simulating NR shows no benefit to the PCr/ATP ratio while inhibition of β-oxidation with trimetazidine shows an even lower PCr/ATP ratio at rest (Table 2).

#### Case 2

Mitochondrial dysfunction can result from compromised mitochondrial biogenesis which results in reduced mitochondrial density. To evaluate the consequence of reduced functioning mitochondria, we simulated a 20% reduction in the volume of active mitochondria present in the cell. The results shown in Fig. 5 suggest that this change significantly reduces energetic reserves indicated by the lower PCr/ATP ratio notably in vigorous exercise conditions. Substrate selection shows a slight preference for carbohydrate oxidation under these conditions. No change in PCr/ATP ratio is observed after simulating NR and trimetazidine at rest and exercise (Table 2).

#### Case 3

A consequence of aging and heart failure is mitochondrial dysfunction resulting from reduced adenine nucleotide metabolites. Simulations with adenine nucleotide pool level similar to that observed for 80-year-old female subjects [44], Fig. 5 shows a steep decline in PCr/ATP ratio compared to the control case at both rest and exercise conditions. In addition, the proposed simulated interventions, including upregulation of PDH to increase carbohydrate oxidation, does little to alter the PCr/ATP ratio (Table 2).

## 4. Discussion

Dysfunction in mitochondrial oxidative metabolism has been suggested to contribute to impaired ATP production of the failing heart [19, 52, 53]. A review by Lopaschuk et al. [54] highlights some of the underlying causes for dysfunction in oxidative mitochondrial metabolism in different etiologies of HF, such as alterations in the redox state of NADH, transcriptional changes of fatty acid oxidative enzymes, and altered fate of glucose. Lopez et al. outlines roles of purine nucleotide metabolism in energetic dysfunction in heart failure [47]. Pharmacological targeting of mitochondrial oxidative metabolism has become an important area of research to address energetic deficiencies in heart failure [55]. Here, we have proposed a quantitative systems pharmacology approach to aid in the analysis of mitochondrial energetic pathways and to identify molecular targets with the potential to ameliorate metabolic inflexibility in HF.

The *in silico* model developed here has several advantages over current state-of-the-art models available in the literature. The first advantage is that the oxidative phosphorylation and TCA cycle portion of the model is constrained by thermodynamics. Previous models including the pathways of interest were fitted using a constraint-based approach which allows a wide range of parameter flexibility to fit steady-state data without physiologically feasible constraints. For example, Moxley et al. used a constraints-based modeling approach to demonstrate how fuel selection in skeletal muscle arises as a function of ATP demand in skeletal muscle [56]. A more mechanistically detailed kinetic model describing substrate selection in the failing heart was developed by Cortassa et al. [27]. The model was used to explore the control of fuel selection by available substrate concentrations and to conduct metabolic control analysis. However, since the model of Cortassa et al. does not capture the role of ATP hydrolysis products in respiratory control it is not intended to reveal insight into the interplay between myocardial energetics and fuel selection. One important advance of the present model is that it includes a model of allosteric regulation of pyruvate dehydrogenase (PDH), which is a crucial regulator of mitochondrial fuel selection. PDH is allosterically inhibited by its products, AcCoA and NADH, and is tightly regulated through pyruvate dehydrogenase kinase (PDK), which phosphorylates PDH to the inactive state. Furthermore, PDK is also regulated by the products of PDH, AcCoA and NADH activate PDK while CoASH, NAD, and pyruvate inhibit the function of PDK. While the model does not explicitly describe PDK, we have incorporated the feedback regulation reflected by the ratios of AcCoA/CoASH, and NADH/NAD, and effect of pyruvate in influencing the Vmax parameter of PDH highlighted in Fig. 2B.

Furthermore, the in silico model developed here shows substrate selection as an emergent property of the *in silico* model resulting from incorporating regulation of PDH and the integration of the β-oxidation pathway as illustrated in Fig. 4A. It has been shown that metabolic substrate selection exhibits preference for carbohydrates in the fed state and with increased ATP demand [57]. In addition to the predicted changes in carbohydrate oxidation with increasing pyruvate, a shift occurs from fatty acid oxidation to carbohydrate oxidation from rest to exercise as shown in Fig. 4B. One of the key questions we sought to address is how substrate selection influences energetics. PCr/ATP ratio is often cited as an energetic biomarker to assess HF in ailing patients [58-60]; therefore, our simulations investigated PCr/ATP ratios with varying ratios of substrate. We found no substantial influence of pyruvate versus fatty acid use on PCr/ATP ratio in simulated healthy cardiomyocytes suggesting cardiomyocytes will use any available fuel to maintain PCr/ATP in dynamic conditions, in agreement with the conclusion from Depre et al. that the heart is broadly able to support normal function in a wide range of availability different substrates [61]. Under conditions where physiological levels of both fatty acid and pyruvate are available, alterations to the Vmax of PDH and citrate synthase are predicted to result in changes to J_PDH_/J_CS_ ratio, but not in PCr/ATP. Exercise intolerance is also a common symptom among HF patients. Increasing the rate of ATP hydrolysis to represent rest and exercise conditions, the model also simulates declining PCr/ATP with increasing exercise level in healthy cardiomyocytes (Fig. 4C). The 30% relative decline agrees with similar declines observed in skeletal muscle during vigorous exercise [62].

Our next objective was to compare substrate selection, energetic measures PCr/ATP, mitochondrial VO_2_, and Pi accumulation of the *in silico* model representing intact myocardium under different mitochondrial dysfunction conditions as described in literature [63]. The first case represents the often cited downregulation of PDH in HF [46, 64]. Case 2 represents a reduction in functional mitochondrial volume per cell [65] and Case 3 has reduced adenine nucleotide pools Pi, PCr, ATP, ADP consistent with aging and advanced heart failure [47, 66] In all of these three cases, simulation results indicate a shift toward increased glucose oxidation, which is predicted to be an adaptive response controlled by feedback mechanisms acting on PDH activity. This finding is consistent with Dodd et al. [14] where the increase in PDH flux in spontaneous hypertensive rat heart is the result of decreases in PDK, the key regulating enzyme of PDH activity [14]. Interestingly, Case 3 (reduced nucleotide pools) was the only case of energetic dysfunction tested that exhibited a reduced PCr/ATP ratio in the rest condition (Fig. 5). The mitochondrial dysfunction in Case 3 is represented by initializing the model with decreased phosphate pools which is reflected by the lower PCr/ATP ratio. The other two cases showed no difference at rest in the PCr/ATP ratio suggesting measurements of PCr/ATP may be in imperfect biomarker to assess HF patients under normal resting conditions. It has been noted by Bottomley et al. [67] and confirmed by Solaiyappan et al. [68] that PCr/ATP measurements alone are not sensitive enough to assess the rate of ATP delivery and the ratio underestimates the quantity of high-energy phosphate metabolites in HF [69].

Conversely, the *in silico* model simulated exercise conditions manifest significant decreases in PCr/ATP ratio for all three cases (Fig 5). These simulations corroborate clinical observations that HF symptoms are exacerbated under stress conditions, such as during exercise.

Finally, we investigated the mechanism of action of several interventions hypothesized to treat HF. To summarize some of our key findings, DCA, a promoter of PHD flux, had the greatest impact on PCr/ATP ratio in case 1, where PDH is initially downregulated and had no bearing on the other two cases of dysfunction (Table 2). We also simulated the direct mitochondrial effects of nicotinamide riboside (NR) which improves the efficiency of the electron transport chain since NAD and NADH play a crucial role in substrate selection. It has been postulated that the imbalance of NAD/NADH ratio is implicated in the development and progression of HF [70]. To test the hypothesis that altered NAD/NADH pools in the mitochondria can alter energetics, we simulated NR supplementation by increasing total NAD^+^ pools by 50%. No significant changes were observed in PCr/ATP or MVO_2_ for all cases of mitochondrial dysfunction investigated (Table 2). The function of NAD pools in HF energetics may play a bigger role in other instances of mitochondrial dysfunction which have not been investigated here. In addition, the benefits of NR in HF may stem from downstream effects of sirtuin signaling, such as mitochondrial biogenesis which was not accounted for in this simulation. Lastly, we simulated the effects of trimetazidine, a potent inhibitor of mckat enzyme in β-oxidation. One hypothesis is that by inhibiting fatty acid oxidation, the heart can utilize glucose as a more efficient fuel for ATP production [71]. In the cases of mitochondrial dysfunction tested here, we did not observe any benefit in ATP production by inhibiting β-oxidation, the PCr/ATP ratio was either not different from no treatment or less than no treatment (Table 2). Here we propose the most appropriate treatment to mitigate alterations in PCr/ATP may depend on etiology of HF. Expanding on this work to further explore the multifactorial causes of metabolic inflexibility in HF may also provide additional insights into metabolic targets for HF.

## 5. Limitations

The model we have presented has been validated against disparate data sets under different conditions and across several species which increases our confidence that model effectively captures the energetic phenotype and, at least partially, the mechanism underlying it. One of the simplifying assumptions of our modeling effort is that the model was developed irrespective to species differences that may be present due to the limited amount of data from humans. The parameters were fitted using *in vitro* data from isolated mouse and rat mitochondria while the model describing intact myocardium incorporates data from dogs and humans. We assume the mitochondrial machinery as well as mechanisms of regulation are conserved across species which allows us to extend our predictions from preclinical species to humans.

Any scientific model, mathematical or otherwise, is a simplification and incomplete, and the field of model-informed drug development (MIDD) [72] highlights the context-of-use as one of the key assessment elements of model evaluation and applicability assessment. The context-of-use of this model while limited to exploration of the underlying mechanisms, provides a step forward to previous work and is an additional tool for generating investigative hypotheses of underlying feature of mitochondrial contribution to energetic phenotypes of HF patients that can be further tested in in vitro, in vivo and/or clinical settings through analyses of collected data. Limitations of the current work include the lack of compensatory mechanisms via metabolic remodeling; the perturbations representing HF are employed as acute effects. Further iterations of the computational models representing *in vitro* mitochondria and intact myocardium would benefit from including more detailed regulation on glucose and lactate to evaluate the possible energetic ramifications of uncoupling glycolysis from glucose oxidation as described by Fillmore et al. [73]. In addition, mitochondrial anaplerosis pathways could be incorporated, such as ketone bodies, branched chain amino acids, and pyruvate carboxylase.

The *in silico* intact myocardium model calibration and validation data is most closely related to HFrEF; however, some of the mechanisms may be relevant for HF with preserved ejection fraction (HFpEF). As more data from HFpEF studies becomes available, the *in silico* outputs should be revalidated for this subset of HF patients.

## 6. Conclusions

To the best of our knowledge, this is the first quantitative investigation into cardiac energetics of HF subgroups with and without an intervention at rest and exercise. We first established the *in silico* model of *in vitro* scenarios reproduces key biological experiments in isolated mitochondria which serve as the foundation for a mechanistic understanding of intact myocardial kinetics. We then found the *in silico* model recapitulates intact myocardium substrate selection in fed and fasted conditions and under different levels of workload. Different model simulated scenarios representing mitochondrial HF show that different mitochondrial HF phenotypes can cause marked changes in energetic measures that may also differ from rest to exercise. Improvements in energetic measures depend on both the mitochondrial HF phenotype and the intervention employed. The simulation studies suggest that identifying HF phenotype subpopulations is an important factor for the development of a successful therapeutic intervention.

