## Appendix for "Development of a Quantitative Systems Pharmacology Model to Interrogate Mitochondrial Metabolism in Heart Failure"

**Appendix: Model Formulation and Description of Codes**

Overview

Computer codes to simulate the coupled pathways of the tricarboxylic acids cycle and beta oxidation were created to meet the following objectives:

- To evaluate substrate selection (pyruvate vs. fatty acids) in mitochondria in cardiac muscle under various conditions, such as during exercise versus rest
- To identify molecular targets within energetic pathways of the mitochondria to augment metabolic inflexibility by improving energetic reserves

The complete model structure is illustrated in Fig. 2. The model takes the form of 107 ordinary differential equations, based on integrating simulation modules from Wu et al. [25] for the TCA cycle and van Eunen et al.[20] for beta oxidation. At total of eight parameters associated with the beta oxidation component were adjusted to match data obtained from respiration of rat cardiac mitochondria fueled by acyl-carnatines of various chain lengths. The identified model was validated based on data from isolated mouse cardiac mitochondria from Fisher-Wellman et al. [42].

Model Development

Integrating the existing models of β-oxidation model from van Eunen et al. [20] and the oxidative phosphorylation model from Wu et al. [25] required several adjustments. To reconcile the units between the two models as the β-oxidation model reports V_max_ values in μmol/min/mg protein and the oxidative phosphorylation model has units of mol/L/sec. Using the provided value for L mito/ mg protein from van Eunen et al. and converting minutes to seconds, results in both models operating with units of mol/L/sec for fluxes. Furthermore, rates of change of metabolic concentrations were updated to account for the combined set of flux expressions from both simulation modules. The operations of the two modules are linked through shared species, including NAD, NADH, CoA, and acetyl-CoA. Direct interactions between these modules are through three conserved mitochondrial metabolite pools with species that participate in reactions in both modules. These three conserved pools are total coenzyme-A , (ie. free coenzyme-A, acetyl-CoA, acyl-CoA species), the total NAD + NADH pool and the total FAD + FADH2 pool.

Parameters for PDH and citrate synthase flux were adjusted to replicate observations on substrate selection. PDH regulation is incorporated in the model by a function representing phosphorylation and dephosphorylation by Pyruvate dehydrogenase kinase. The V_max_ of PDH is altered by a factor that varies by the ratios of acetyl-coA to CoASH, NADH to NAD^+^, ATP to ADP, and ~1/Pyruvate present in the mitochondrial matrix as shown below in Eqs. C4-C6.

Key Assumptions and Simplifications

- The model does not account for changes in concentrations of cytosolic carnitine, coenzyme A, malonyl-CoA and matrix carnitine and, thus, concentrations of these species are clamped at constant values.
- Cellular biochemical processes contributing to ATP hydrolysis are lumped in to single ATP hydrolysis flux. The rate of this flux is varied from rest (0.05 mM/s) to exercise (1.5 mM/s) to simulate the changes in the demand for ATP.
- The pH of the matrix, K^+^, and Mg^+^ was fixed. This simplication was made to reduce the complexity of merging the two models.

Simulation Conditions

For simulations of in vitro experiments using suspensions of isolated mitochondria, concentrations of extramitochondrial substrates introduced to the respiration media vary as governed by exchange between buffer and mitochondrial compartments. For simulation of intact myocardial metabolism, cytosolic primary substrates pyruvate and palmitoyl-coA are clamped at concentration values choses to represent fasting and fed states, as detailed below.

**Table A1:** State Variable Definitions

| **Variables** | **Definition** | **Units** | **Matlab Code Representation** |
| --- | --- | --- | --- |
| ΔΨ | Mitochondrial membrane potential | mV | dPsi |
| $[H^{+}]$ | Concentration of H^+^ ion | mol (L water)^-1^ | H |
| $[K^{+}]$ | Concentration of K^+^ ion | mol (L water)^-1^ | K |
| $[{Mg}^{+2}]$ | Concentration of Mg^2+^ ion | mol (L water)^-1^ | Mg |
| $[NADH]$ | Concentration of NADH | mol (L water)^-1^ | NADH |
| $[NAD]$ | Concentration of NAD | mol (L water)^-1^ | NAD |
| $[QH_{2}]$ | Concentration of reduced ubiquinol | mol (L water)^-1^ | QH2 |
| $[COQ]$ | Concentration of oxidized ubiquinol | mol (L water)^-1^ | COQ |
| $[ATP]$ | Concentration of ATP | mol (L water)^-1^ | ATP |
| $[ADP]$ | Concentration of ADP | mol (L water)^-1^ | ADP |
| $[GTP]$ | Concentration of GTP | mol (L water)^-1^ | GTP |
| $[GDP]$ | Concentration of GDP | mol (L water)^-1^ | GDP |
| $[PI]$ | Concentration of inorganic phosphate | mol (L water)^-1^ | PI |
| $[PYR]$ | Concentration of pyruvate | mol (L water)^-1^ | PYR |
| $[COASH]$ | Concentration of CoA-SH | mol (L water)^-1^ | CoAMAT |
| $[ACCOA]$ | Concentration of acetyl-CoA | mol (L water)^-1^ | ACCOA |
| $[OAA]$ | Concentration of oxaloacetate | mol (L water)^-1^ | OAA |
| $[CIT]$ | Concentration of citrate | mol (L water)^-1^ | CIT |
| $[ICIT]$ | Concentration of isocitrate | mol (L water)^-1^ | ICIT |
| $[AKG]$ | Concentration of α-ketoglutarate | mol (L water)^-1^ | AKG |
| $[SCOA]$ | Concentration of succinyl-CoA | mol (L water)^-1^ | SCOA |
| $[SUC]$ | Concentration of succinate | mol (L water)^-1^ | SUC |
| $[FUM]$ | Concentration of fumarate | mol (L water)^-1^ | FUM |
| $[MAL]$ | Concentration of malate | mol (L water)^-1^ | MAL |
| $[ASP]$ | Concentration of aspartate | mol (L water)^-1^ | ASP |
| $[GLU]$ | Concentration of glutamate | mol (L water)^-1^ | GLU |
| $[O_{2}]$ | Concentration of oxygen | mol (L water)^-1^ | PO2 |
| $[{CO}_{2}tot]$ | Concentration of total carbon dioxide | mol (L water)^-1^ | CO2tot |
| $[C_{n}\text{acyclcarn}\text{]}$ | Concentration of C_n_ acyl-carnitine (n=4,6,8,10,12,14, or 16) | mol (L water)^-1^ | C16Carn_cy |
| ${[C}_{n}AcylCoA]$ | Concentration of C_n_ Acyl-CoA (n=4,6,8,10,12,14, or 16) | mol (L water)^-1^ | C16CoA_m |
| ${[C}_{n}EnoylCoA]$ | Concentration of C_n_ Enoyl-CoA (n=4,6,8,10,12,14, or 16) | mol (L water)^-1^ | C16EnoylCoA_m |
| ${[C}_{n}HCoA]$ | Concentration of C_n_ Hydroxyacyl-CoA (n=4,6,8,10,12,14, or 16) | mol (L water)^-1^ | C16OHCoA_m |
| ${[C}_{n}KetoCoA]$ | Concentration of C_n_ Ketoacyl-CoA (n=4,6,8,10,12,14, or 16) | mol (L water)^-1^ | C16KetoCoA_m |
| $[GLC]$ | Concentration of glucose | mol (L water)^-1^ | GLC |
| $[G6P]$ | Concentration of glucose-6-phosphate | mol (L water)^-1^ | G6P |
| $[PCr]$ | Concentration of phosphate creatine | mol (L water)^-1^ | PCr |
| $[Cr]$ | Concentraiton of creatine | mol (L water)^-1^ | Cr |
| $[Cred]$ | Concentration of reduced cytochrome c | mol (L water)^-1^ | Cred |
| ${[\_\_]}_{x}$ | Species concentration in mitochondria matrix | mol (L water)^-1^ | _x |
| ${[\_\_]}_{i}$ | Species concentration in mitochondrial inter-mebrane space | mol (L water)^-1^ | _i |
| ${[\_\_]}_{c}$ | Species concentration cytosol or buffer | mol (L water)^-1^ | _c |

**Table A2:** Flux Definitions

| **Flux Variable** | **Definition** | **Units** | **Matlab Code Representation** |
| --- | --- | --- | --- |
| $J_{C1}$ | Complex I | mol s^-1^ (L mito)^-1^ | J_C1 |
| $J_{C3}$ | Complex III | mol s^-1^ (L mito)^-1^ | J_C3 |
| $J_{C4}$ | Complex IV | mol s^-1^ (L mito)^-1^ | J_C4 |
| $J_{F1}$ | F_1_F_0_ ATPase reaction | mol s^-1^ (L mito)^-1^ | J_F1 |
| $J_{ANT}$ | Adenine nucleotide translocase | mol s^-1^ (L mito)^-1^ | J_ANT |
| $J_{Hle}$ | Proton Leak | mol s^-1^ (L mito)^-1^ | J_Hle |
| $J_{ASPGLU}$ | Aspartate – Glutamate antiporter | mol s^-1^ (L mito)^-1^ | J_ASP_GLU |
| $J_{ndk}$ | Nucleoside diphosphokinase | mol s^-1^ (L mito)^-1^ | J_ndk |
| $J_{scoas}$ | Succinyl-CoA synthetase | mol s^-1^ (L mito)^-1^ | J_scoas |
| $J_{PIHt}$ | Phosphate-hydrogen co-transporter | mol s^-1^ (L mito)^-1^ | J_PI1 |
| $J_{MALPI}$ | Malate/phosphate antiporter | mol s^-1^ (L mito)^-1^ | J_MAL_PI |
| $J_{pdh}$ | Pyruvate dehydrogenase | mol s^-1^ (L mito)^-1^ | J_pdh |
| $J_{isod}$ | Isocitrate dehydrogenase | mol s^-1^ (L mito)^-1^ | J_isod |
| $J_{mdh}$ | Malate dehydrogenase | mol s^-1^ (L mito)^-1^ | J_mdh |
| $J_{boxNADH}$ | NADH from β-oxidation | mol s^-1^ (L mito)^-1^ | J_boxnadh |
| $J_{sdh}$ | Succinate dehydrogenase | mol s^-1^ (L mito)^-1^ | Jsdh |
| $J_{PYRH}$ | Pyruvate – H^+^ co-transporter | mol s^-1^ (L mito)^-1^ | J_PYR_H |
| $J_{cits}$ | Citrate synthetase | mol s^-1^ (L mito)^-1^ | J_cits |
| $J_{boxACCoA}$ | Acetyl-CoA from β-oxidation | mol s^-1^ (L mito)^-1^ | J_boxaccoa |
| $J_{acon}$ | Aconitase | mol s^-1^ (L mito)^-1^ | J_acon |
| $J_{CITMAL}$ | Citrate/malate antiporter | mol s^-1^ (L mito)^-1^ | J_CIT_MAL |
| $J_{got}$ | Glutamate oxaloacetate transaminase | mol s^-1^ (L mito)^-1^ | J_got |
| $J_{AKGMAL}$ | α-Ketoglutarate/malate antiporter | mol s^-1^ (L mito)^-1^ | J_AKG_MAL |
| $J_{SUCMAL}$ | Succinate/malate antiporter | mol s^-1^ (L mito)^-1^ | J_SUC_MAL |
| $J_{fum}$ | Fumarase | mol s^-1^ (L mito)^-1^ | J_fum |
| $J_{GLUH}$ | Glutamate – H^+^ co-transporter | mol s^-1^ (L mito)^-1^ | J_GLU_H |
| $J_{ASPGLU}$ | Aspartate/glutamate antiporter | mol s^-1^ (L mito)^-1^ | J_ASP_GLU |
| $J_{cactCn}$ | Carnitine-acyl-carnitine translocase | mol s^-1^ (L mito)^-1^ | reaction_vcactC10 |
| $J_{cpt2Cn}$ | Carnitine-palmitoyl transferase 2 | mol s^-1^ (L mito)^-1^ | reaction_vcpt2C12 |
| $J_{vlcadCn}$ | Very-long-chain acyl-CoA dehydrogenase | mol s^-1^ (L mito)^-1^ | reaction_vvlcadC16 |
| $J_{lcadCn}$ | Long-chain acyl-CoA dehydrogenase | mol s^-1^ (L mito)^-1^ | reaction_vlcadC16 |
| $J_{crotCn}$ | Crotonase | mol s^-1^ (L mito)^-1^ | reaction_vcrotC12 |
| $J_{mtpCn}$ | Mitochondrial trifunctional protein | mol s^-1^ (L mito)^-1^ | reaction_vmtpC14 |
| $J_{mschadCn}$ | Medium/short-chain hydroxyacyl-CoA dehydrogenase | mol s^-1^ (L mito)^-1^ | reaction_vmschadC4 |
| $J_{mckatCn}$ | Medium-chain ketoacyl-CoA thiolase | mol s^-1^ (L mito)^-1^ | reaction_vmckatC4 |
| $J_{mcadCn}$ | Medium-chain acyl-CoA dehydrogenase | mol s^-1^ (L mito)^-1^ | reaction_vmcadC4 |
| $J_{scadCn}$ | Short-chain acyl-CoA dehydrogenase | mol s^-1^ (L mito)^-1^ | reaction_vscadC4 |
| $J_{cpt1Cn}$ | Carnitine-palmitoyl transferase 1 | mol s^-1^ (L mito)^-1^ | reaction_vcpt1C16 |
| $J_{FADHsink}$ | FADH sink | mol s^-1^ (L mito)^-1^ | reaction_vfadhsink |
| $J_{AKi}$ | Mitochondrial adenylate kinase | mol s^-1^ (L mito)^-1^ | J_Aki |
| $J_{PIt}$ | Phosphate transport across outer membrane | mol s^-1^ (L mito)^-1^ | J_PI2 |
| $J_{GLUt}$ | Glutamate transport across outer membrane | mol s^-1^ (L mito)^-1^ | J_GLUt |
| $J_{PYRt}$ | Pyruvate transport across outer membrane | mol s^-1^ (L mito)^-1^ | J_PYRt |
| $J_{CITt}$ | Citrate transport across outer membrane | mol s^-1^ (L mito)^-1^ | J_CITt |
| $J_{ICITt}$ | Isocitrate transport across outer membrane | mol s^-1^ (L mito)^-1^ | J_ICITt |
| $J_{AMPt}$ | AMP transport across outer membrane | mol s^-1^ (L mito)^-1^ | J_AMP |
| $J_{AKGt}$ | α-Ketoglutarate transport across outer membrane | mol s^-1^ (L mito)^-1^ | J_AKGt |
| $J_{SUCt}$ | Succinate transport across outer membrane | mol s^-1^ (L mito)^-1^ | J_SUCt |
| $J_{MALt}$ | Malate transport across outer membrane | mol s^-1^ (L mito)^-1^ | J_MALt |
| $J_{ASPt}$ | Aspartate transport across outer membrane | mol s^-1^ (L mito)^-1^ | J_ASPt |
| $J_{ATPt}$ | ATP transport across outer membrane | mol s^-1^ (L mito)^-1^ | J_ATP |
| $J_{ADPt}$ | ADP transport across outer membrane | mol s^-1^ (L mito)^-1^ | J_ADP |
| $J_{HK}$ | Hexokinase | mol s^-1^ (L buffer)^-1^ | J_HK |
| $J_{AtC}$ | ATP Hydrolysis | mol s^-1^ (L cytosol)^-1^ | J_AtC |
| $J_{CKc}$ | Creatine Kinase | mol s^-1^ (L cytosol)^-1^ | J_CKe |

**Table A3:** Parameter Definitions

| **Parameter** | **Definition** | **Value** | **Units** | **Matlab Code Representation** |
| --- | --- | --- | --- | --- |
| $W_{x}$ | Matrix water space fraction | 0.6514 | L mito (L cell)^-1^ | W_x |
| $V_{x}$ | Volume of matrix per mg protein | 1.8 x 10^-6^ | L mito (mg protein) ^-1^ | compartment_VMAT |
| $C_{IM}$ | Capacitance of the inner membrane | 6.75 x 10^-6^ | mol L^-1^ mV^-1^ | Fixed constant |
| $n_{A}$ | H+ stoichiometry coefficient for F1F0-ATPase | 3 | unitless | n_A |
| $W_{i}$ | Inter-membrane space water fraction | 0.0724 | L mito (L cell)^-1^ | W_i |
| $W_{c}$ | Cytoplasm water fraction | 0.843 | L cyto (L cell)^-1^ | W_c |
| $V_{mito}$ | Mitochondria Volume | 0.288 | L mito (L cell)^-1^ | Vmito |
| $V_{cyto}$ | Cytoplasm Volume | 0.680 | L cyto (L cell)^-1^ | Vcyto |
| ${NAD}_{tot}$ | Total matrix NAD(H) concentration | 2.97 x 10^-3^ | mol (L matrix)^-1^ | NADtot |
| $Q_{tot}$ | Total matrix ubiquinol concentration | 1.35 x 10^-3^ | mol (L matrix)^-1^ | Qtot |
| ${cytC}_{tot}$ | Total inter-membrane (IM) cytochrome c concentration | 2.70 x 10^-3^ | mol (L IM)^-1^ | Ctot |
| ${CoA}_{tot}$ | Total Coenzyme A | 11 x 10^-3^ | L mito (L cell)^-1^ | Const_species_CoAMATt |
| $V_{maxPDH}$ | Final Vmax of PDH | calculated | mol s^-1^ (L mito)^-1^ | aa*Vmf_PDH |
| ${PDH}_{cf}$ | PDH correction factor | calculated | unitless | aa |
| ${Vmf}_{PDH}$ | PDH enzyme activity | 2.0 x 10^-3^ | mol s^-1^ (L mito)^-1^ | Vmf_PDH |
| $ra$ | Ratios of PDH regulators | calculated | unitless | rat |
| $ref\_PYR$ | Pyruvate proportionality constant | 1.94 x 10^-3^ | M | ref_pyr |

Governing Equations

The model governing differential equations are are listed below organized under subheadings for variables representing membrane potential, mitochondrial matrix variables (denoted by subscript x), intermembrane space variables(denoted by subscript i), cytosolic or external space variables (denoted by subscript c), and PDH regulation. The external space variables used depend on the type of simulation considered (*ex vivo* isolated mitochondria or intact myocardium system.) Detailed descriptions for each flux (J_species_ ) expression are found in the appendices of Wu et. al [25] and van Eunen et al. [20]. Equations are labeled numerically with prefix A, B or C, representing the origin of the equation if from the model of oxidative phosphorylation, β-oxidation, or specific to the combined model, respectively.

Mitochondrial Inner Membrane Electrical Potential:

$C_{\text{ΙΜ}}\frac{d\Delta\Psi}{dt}=+4J_{C1}+2J_{\text{C3}}+4J_{\text{C4}}-n_{A}J_{\text{F1}}-J_{\text{ANT}}-J_{\text{Hle}}+J_{\text{ASPGLU}}$	Eq. A1

Mitochondrial Matrix:

$d[\text{ATP}]_{x}/dt=(+J_{\text{ndk}}+J_{\text{F1}}-J_{\text{ANT}})/W_{x}$	Eq. A2

$d[\text{ADP}]_{x}/dt=(-J_{\text{ndk}}-J_{\text{F1}}+J_{\text{ANT}})/W_{x}$	Eq. A3

$d[\text{AMP}]_{x}/dt=0/W_{x}$	Eq. A4

$d[\text{GTP}]_{x}/dt=(+J_{\text{scoas}}-J_{\text{ndk}})/W_{x}$	Eq. A5

$d[\text{GDP}]_{x}/dt=(-J_{\text{scoas}}+J_{\text{ndk}})/W_{x}$	Eq. A6

$d[\text{PI}]_{x}/dt=(-J_{\text{scoas}}-J_{\text{F1}}+J_{\text{PIHt}}-J_{\text{MALPI}})/W_{x}$	Eq. A7

$d[\text{NADH}]_{x}/dt=(+J_{\text{pdh}}+J_{\text{isod}}+J_{\text{akgd}}+J_{\text{mdh}}-J_{\text{C1}})/W_{x}$ + $J_{\text{boxNADH}}$	Eq. C1

$d[\text{Q}\text{H}_{2}]_{x}/dt=(+J_{\text{sdh}}+J_{\text{C1}}-J_{\text{C3}})/W_{x}$	Eq. A9

$d[\text{PYR}]_{x}/dt=(-J_{\text{pdh}}+J_{\text{PYRH}})/W_{x}$	Eq. A10

$d[\text{ACCOA}]_{x}/dt=(-J_{\text{cits}}+J_{\text{pdh}})/W_{x}$ ${+J}_{\text{boxACCoA}}$	Eq. C2

$d[\text{CIT}]_{x}/dt=(+J_{\text{cits}}-J_{\text{acon}}+J_{\text{CITMAL}})/W_{x}$	Eq. A12

$d[\text{ICIT}]_{x}/dt=(+J_{\text{acon}}-J_{\text{isod}})/W_{x}$	Eq. A13

$d[\text{AKG}]_{x}dt=(+J_{\text{isod}}-J_{\text{akgd}}-J_{\text{got}}+J_{\text{AKGMAL}})/W_{x}$	Eq. A14

$d[\text{SCOA}]_{x}/dt=(+J_{\text{akgd}}-J_{\text{scoas}})/W_{x}$	Eq. A15

$d[\text{SUC}]_{x}/dt=(+J_{\text{scoas}}-J_{\text{sdh}}+J_{\text{SUCMAL}})/W_{x}$	Eq. A17

$d[\text{FUM}]_{x}/dt=(+J_{\text{sdh}}-J_{\text{fum}})/W_{x}$	Eq. A18

$d[\text{MAL}]_{x}/dt=(+J_{\text{fum}}-J_{\text{mdh}}+J_{\text{MALPI}}-J_{\text{AKGMAL}}-J_{\text{CITMAL}}-J_{\text{SUCMAL}})/W_{x}$	Eq. A19

$d[\text{OAA}]_{x}/dt=(-J_{\text{cits}}+J_{\text{mdh}}+J_{\text{got}})/W_{x}$	Eq. A20

$d[\text{GLU}]_{x}/dt=(+J_{\text{got}}+J_{\text{GLUH}}-J_{\text{ASPGLU}})/W_{x}$	Eq. A21

$d[\text{ASP}]_{x}/dt=(-J_{\text{got}}+J_{\text{ASPGLU}})/W_{x}$	Eq. A22

$d[O_{2}]_{x}/dt=0/W_{x}$	Eq. A23

$d[\text{C}\text{O}_{2}\text{tot}]_{x}/dt=0/W_{x}$	Eq. A24

$d[C_{16}\text{Ac}{ylcarn]}_{x}/dt=(+J_{\text{cactC16}}-J_{\text{cpt2C16}})/V_{x}$	Eq. B2

$d[C_{16}\text{AcylCoA}]_{x}/dt=(+J_{\text{cpt2C16}}-J_{\text{vlcadC16}}-J_{\text{lcadC16}})/V_{x}$	Eq. B3

$d[C_{16}\text{EnoylCoA}]_{x}/dt=(J_{\text{vlcadC16}}+J_{\text{lcadC16}}-J_{\text{crotC16}}-J_{\text{mtpC16}})/V_{x}$	Eq. B4

$d[C_{16}\text{H}{CoA]}_{x}/dt=(+J_{\text{crotC16}}-J_{\text{mschadC16}})/V_{x}$	Eq. B5

$d[C_{16}\text{Keto}{CoA]}_{x}/dt=(+J_{\text{mschadC16}}-J_{\text{mckatC16}})/V_{x}$	Eq. B6

$d[C_{14}\text{Ac}{ylcarn]}_{x}/dt=(+J_{\text{cactC14}}-J_{\text{cpt2C14}})/V_{x}$	Eq. B8

$d[C_{14}\text{AcylCoA}]_{x}/dt=(+J_{\text{cpt2C14}}+J_{\text{mtpC16}}+J_{\text{mckatC16}}-J_{\text{vlcadC14}}-J_{\text{lcadC14}})/V_{x}$	Eq. B9

$d[C_{14}\text{EnoylCoA}]_{x}/dt=(J_{\text{vlcadC14}}+J_{\text{lcadC14}}-J_{\text{crotC14}}-J_{\text{mtpC14}})/V_{x}$	Eq. B10

$d[C_{14}\text{H}{CoA]}_{x}/dt=(+J_{\text{crotC14}}-J_{\text{mschadC14}})/V_{x}$	Eq. B11

$d[C_{14}\text{Keto}{CoA]}_{x}/dt=(+J_{\text{mschadC14}}-J_{\text{mckatC14}})/V_{x}$	Eq. B12

$d[C_{12}\text{Ac}{ylcarn]}_{x}/dt=(+J_{\text{cactC12}}-J_{\text{cpt2C12}})/V_{x}$	Eq. B14

$d[C_{12}\text{AcylCoA}]_{x}/dt=(+J_{\text{cpt2C12}}+J_{\text{mtpC14}}+J_{\text{mckatC14}}-J_{\text{vlcadC12}}-J_{\text{lcadC12}}-J_{\text{mcadC12}})/V_{x}$	Eq. B15

$d[C_{12}\text{EnoylCoA}]_{x}/dt=(-J_{\text{vlcadC12}}+J_{\text{lcadC12}}+J_{\text{mcadC12}}-J_{\text{crotC12}}-J_{\text{mtpC12}})/V_{x}$	Eq. B16

$d[C_{12}\text{H}{CoA]}_{x}/dt=(+J_{\text{crotC12}}-J_{\text{mschad2C12}})/V_{x}$	Eq. B17

$d[C_{12}\text{Keto}{CoA]}_{x}/dt=(+J_{\text{mschadC12}}-J_{\text{mckat2C12}})/V_{x}$	Eq. B18

$d[C_{10}\text{Ac}{ylcarn]}_{x}/dt=(+J_{\text{cactC10}}-J_{\text{cpt2C10}})/V_{x}$	Eq. B20

$d[C_{10}\text{AcylCoA}]_{x}/dt=(+J_{\text{cpt2C10}}+J_{\text{mtpC12}}+J_{\text{mckatC12}}-J_{\text{lcadC10}}-J_{\text{mcadC10}})/V_{x}$	Eq. B21

$d[C_{10}\text{EnoylCoA}]_{x}/dt=(+J_{\text{lcadC10}}+J_{\text{mcadC10}}-J_{\text{crotC10}}-J_{\text{mtpC10}})/V_{x}$	Eq. B22

$d[C_{10}\text{H}{CoA]}_{x}/dt=(+J_{\text{crotC10}}-J_{\text{mschadC10}})/V_{x}$	Eq. B23

$d[C_{10}\text{Keto}{CoA]}_{x}/dt=(+J_{\text{mschadC10}}-J_{\text{mckatC10}})/V_{x}$	Eq. B24

$d[C_{8}\text{Ac}{ylcarn]}_{x}/dt=(+J_{\text{cactC8}}-J_{\text{cpt2C8}})/V_{x}$	Eq. B26

$d[C_{8}\text{AcylCoA}]_{x}/dt=(+J_{\text{cpt2C8}}+J_{\text{mtpC10}}+J_{\text{mckatC10}}-J_{\text{lcadC8}}-J_{\text{mcadC8}})/V_{x}$	Eq. B27

$d[C_{8}\text{EnoylCoA}]_{x}/dt=(+J_{\text{lcadC8}}+J_{\text{mcadC8}}-J_{\text{crotC8}}-J_{\text{mtpC8}})/V_{x}$	Eq. B28

$d[C_{8}\text{H}{CoA]}_{x}/dt=(+J_{\text{crotC8}}-J_{\text{mschadC8}})/V_{x}$	Eq. B29

$d[C_{8}\text{Keto}{CoA]}_{x}/dt=(+J_{\text{mschadC8}}-J_{\text{mckatC8}})/V_{x}$	Eq. B30

$d[C_{6}\text{Ac}{ylcarn]}_{x}/dt=(+J_{\text{cactC6}}-J_{\text{cpt2C6}})/V_{x}$	Eq. B32

$d[C_{6}\text{AcylCoA}]_{x}/dt=(+J_{\text{cpt2C6}}+J_{\text{mtpC8}}+J_{\text{mckatC8}}-J_{\text{mcadC6}}-J_{\text{scadC6}})/V_{x}$	Eq. B33

$d[C_{6}\text{EnoylCoA}]_{x}/dt=(+J_{\text{mcadC6}}+J_{\text{scadC6}}-J_{\text{crotC6}})/V_{x}$	Eq. B34

$d[C_{6}\text{H}{CoA]}_{x}/dt=(+J_{\text{crotC6}}-J_{\text{mschadC6}})/V_{x}$	Eq. B35

$d[C_{6}\text{Keto}{CoA]}_{x}/dt=(+J_{\text{mschadC6}}-J_{\text{mckatC6}})/V_{x}$	Eq. B36

$d[C_{4}\text{Ac}{ylcarn]}_{x}/dt=(+J_{\text{cactC4}}-J_{\text{cpt2C4}})/V_{x}$	Eq. B38

$d[C_{4}\text{AcylCoA}]_{x}/dt=(+J_{\text{cpt2C4}}+J_{\text{mckatC6}}-J_{\text{mcadC4}}-J_{\text{scadC4}})/V_{x}$	Eq. B39

$d[C_{4}\text{EnoylCoA}]_{x}/dt=(+J_{\text{mcadC4}}+J_{\text{scadC4}}-J_{\text{crotC4}})/V_{x}$	Eq. B40

$d[C_{4}\text{H}{CoA]}_{x}/dt=(+J_{\text{crotC4}}-J_{\text{mschadC4}})/V_{x}$	Eq. B41

$d[C_{4}\text{Keto}{CoA]}_{x}/dt=(+J_{\text{mschadC4}}-J_{\text{mckatC4}})/V_{x}$	Eq. B42

$d[FAD{H]}_{x}/dt=(\sum J_{\text{vlcadCn}}+\sum J_{\text{lcadCn}}+\sum J_{\text{mcadCn}} +\sum J_{\text{vscadCn}}-J_{\text{FADHsink}} )/W_{x}$	Eq. B44

Mitochondrial Inter-Membrane Space:

$d[\text{Cred}]_{i}/dt=(+2J_{\text{C3}}-2J_{\text{C4}})/W_{i}$	Eq. A25

$d[\text{ATP}]_{i}/dt=(+J_{\text{ATP}}+J_{\text{ANT}}+J_{\text{AKi}})/W_{i}$	Eq. A26

$d[\text{ADP}]_{i}/dt=(+J_{\text{ADP}}-J_{\text{ANT}}-2J_{\text{AKi}})/W_{i}$	Eq. A27

$d[\text{AMP}]_{i}/dt=(J_{\text{AMPt}}+J_{\text{AKi}})/W_{i}$	Eq. A28

$d[\text{PI}]_{i}/dt=(-J_{\text{PIHt}}+J_{\text{PIt}}+J_{\text{MALPI}})/W_{i}$	Eq. A29

$d[\text{PYR}]_{i}/dt=(-J_{\text{PYRH}}+J_{\text{PYRt}})/W_{i}$	Eq. A30

$d[\text{CIT}]_{i}/dt=(-J_{\text{CITMAL}}+J_{\text{CITt}})/W_{i}$	Eq. A31

$d[\text{ICIT}]_{i}/dt=(+J_{\text{ICITt}})/W_{i}$	Eq. A32

$d[\text{AKG}]_{i}/dt=(-J_{\text{AKGMAL}}+J_{\text{AKGt}})/W_{i}$	Eq. A33

$d[\text{SUC}]_{i}/dt=(+J_{\text{SUCt}}-J_{\text{SUCMAL}})/W_{i}$	Eq. A34

$d[\text{FUM}]_{i}/dt=0/W_{i}$	Eq. A35

$d[\text{MAL}]_{i}/dt=(-J_{\text{MALPI}}+J_{\text{MALt}}+J_{\text{AKGMAL}}+J_{\text{CITMAL}}+J_{\text{SUCMAL}})/W_{i}$	Eq. A36

$d[\text{GLU}]_{i}/dt=(-J_{\text{GLUH}}+J_{\text{ASPGLU}}+J_{\text{GLUt}})/W_{i}$	Eq. A37

$d[\text{ASP}]_{i}/dt=(-J_{\text{ASPGLU}}+J_{\text{ASPt}})/W_{i}$.	Eq. A38

External Space, *Ex Vivo* Isolated Mitochondria:

$d[\text{ATP}]_{c}/dt=(-J_{\text{ATPt}}-J_{\text{HK}}(1+W_{c}))/W_{c}$	Eq. A39

$d[\text{ADP}]_{c}/dt=(-J_{\text{ADPt}}+J_{\text{HK}}(1+W_{c}))/W_{c}$	Eq. A40

$d[\text{AMP}]_{c}/dt=0/W_{c}$	Eq. A41

$d[\text{PI}]_{c}/dt=-J_{\text{PIt}}/W_{c}$	Eq. A42

$d[\text{PYR}]_{c}/dt=-J_{\text{PYRt}}/W_{c}$	Eq. A43 $d[\text{CIT}]_{c}/dt=-J_{\text{CITt}}/W_{c}$	Eq. A44

$d[\text{ICIT}]_{c}/dt=-J_{\text{ICITt}}/W_{c}$	Eq. A45

$d[\text{AKG}]_{c}/dt=-J_{\text{AKGt}}/W_{c}$	Eq. A46

$d[\text{SUC}]_{c}/dt=-J_{\text{SUCt}}/W_{c}$	Eq. A47

$d[\text{FUM}]_{c}/dt=0/W_{c}$	Eq. A48

$d[\text{MAL}]_{c}/dt=-J_{\text{MALt}}/W_{c}$	Eq. A49

$d[\text{GLU}]_{c}/dt=-J_{\text{GLUt}}/W_{c}$	Eq. A50

$d[\text{ASP}]_{c}/dt=-J_{\text{ASPt}}/W_{c}$	Eq. A51

$d[\text{GLC}]_{c}/dt=-J_{\text{HK}}(1+W_{c})/W_{c}$	Eq. A52

$d[\text{G6P}]_{c}/dt=+J_{\text{HK}}(1+W_{c})/W_{c}$	Eq. A53

$d[\text{PCr}]_{c}/dt=0/W_{c}$;	Eq. A54

External Space, Intact Myocardium:

$d[\text{ATP}]_{c}/dt=\left( -\left( \frac{V_{\text{cyto}}}{V_{\text{mito}}} \right)J_{\text{ATPt}}-J_{\text{AtC}}+J_{\text{CK}}+J_{\text{AKc}} \right)/W_{c}$	Eq. A55

$d[\text{ADP}]_{c}/dt=\left( -\left( \frac{V_{\text{cyto}}}{V_{\text{mito}}} \right)J_{\text{ADPt}}+J_{\text{AtC}}-J_{\text{CK}}-2J_{\text{AKc}} \right)/W_{c}$	Eq. A56

$d[\text{AMP}]_{c}/dt=\left( -\left( \frac{V_{\text{cyto}}}{V_{\text{mito}}} \right)J_{\text{AMPt}}+J_{\text{AKc}} \right)/W_{c}$	Eq. A57

$d[\text{PI}]_{c}/dt=\left( -\left( \frac{V_{\text{cyto}}}{V_{\text{mito}}} \right)J_{\text{PIt}}+J_{\text{AKc}} \right)/W_{c}$	Eq. A58

$d[\text{PYR}]_{c}/dt=-\left( \frac{V_{\text{cyto}}}{V_{\text{mito}}} \right)J_{\text{PYRt}}/W_{c}$	Eq. A59

$d[\text{CIT}]_{c}/dt=-\left( \frac{V_{\text{cyto}}}{V_{\text{mito}}} \right)J_{\text{CITt}}/W_{c}$	Eq. A60

$d[\text{AKG}]_{c}/dt=-\left( \frac{V_{\text{cyto}}}{V_{\text{mito}}} \right)J_{\text{AKGt}}/W_{c}$	Eq. A61

$d[\text{SUC}]_{c}/dt=-\left( \frac{V_{\text{cyto}}}{V_{\text{mito}}} \right)J_{\text{SUCt}}/W_{c}$	Eq. A62

$d[\text{FUM}]_{c}/dt=0/W_{c}$	Eq. A63

$d[\text{MAL}]_{c}/dt=-\left( \frac{V_{\text{cyto}}}{V_{\text{mito}}} \right)J_{\text{MALt}}/W_{c}$	Eq. A64

$d[\text{GLU}]_{c}/dt=-\left( \frac{V_{\text{cyto}}}{V_{\text{mito}}} \right)J_{\text{GLUt}}/W_{c}$	Eq. A65

$d[\text{ASP}]_{c}/dt=-\left( \frac{V_{\text{cyto}}}{V_{\text{mito}}} \right)J_{\text{ASPt}}/W_{c}$	Eq. A66

$d[\text{PCr}]_{c}/dt=-J_{\text{CKc}}/W_{c}$	Eq. A67

$d[C_{16}\text{Ac}{ylCoA]}_{c}/dt=0/W_{c}$	Eq. C3

$d[C_{16}\text{Ac}{ylcarn]}_{c}/dt=(+J_{\text{cpt1C16}}-J_{\text{cactC16}})/W_{c}$	Eq. B1

$d[C_{14}\text{Ac}{ylcarn]}_{c}/dt=-J_{\text{cactC14}}/W_{c}$	Eq. B7

$d[C_{12}\text{Ac}{ylcarn]}_{c}/dt=-J_{\text{cactC12}}/W_{c}$	Eq. B13

$d[C_{10}\text{Ac}{ylcarn]}_{c}/dt=-J_{\text{cactC10}}/W_{c}$	Eq. B19

$d[C_{8}\text{Ac}{ylcarn]}_{c}/dt=-J_{\text{cactC8}}/W_{c}$	Eq. B25

$d[C_{6}\text{Ac}{ylcarn]}_{c}/dt=-J_{\text{cactC6}}/W_{c}$	Eq. B31

$d[C_{4}\text{Ac}{ylcarn]}_{c}/dt=-J_{\text{cactC4}}/W_{c}$	Eq. B37

Assuming constant total concentrations NADtot, Qtot, cytCtot, and Atot for nicotinamide nucleotides, ubiquinol, and cytochrome c, we compute concentrations of the following reactants as:

$[\text{NAD}]_{x}=\text{ NA}\text{D}_{\text{tot}}-[\text{NADH}]_{x}$	Eq. A68

$[\text{COQ}]_{x}=\text{ }\text{Q}_{\text{tot}}-[\text{Q}\text{H}_{2}]_{x}$	Eq. A69

$[\text{Cox}]_{i}=\text{ cyt}\text{C}_{\text{tot}}-[\text{Cred}]_{i}$.	Eq. A70

The coenzyme A pool required the addition of several Co-A species from β-oxidation. Free coenzyme-A (CoASH) is represented by a constant CoA total minus the sum of all other CoA species:

$[\text{CoASH]}=\text{ Constant}\text{CoA}_{\text{tot}}-\sum[\text{CoA}]_{x}$.	Eq. A71

*PDH Regulation*:

PDH is regulated via a phosphorylation-dephosphorylation cycle in which production of the phosphorylated inactive form is stimulated by NADH, ATP, and acyl-coA and dephosphorylation to the activated form is stimulated by pyruvate, NAD, ADP and coenzyme A. We assume that activation and deactivation follow simple Michaelis-Menten kinetics:

$J_{a}=\frac{V_{a}}{1+ {K_{a}}/{(1-a)}}$ and $J_{a}=\frac{V_{i}}{1+ {K_{i}}/a}$

where $J_{a}$ and $J_{i}$ are the activation and inactivation rates, *a* is the fraction of the enzyme in the active form, $V_{a}$ are the Vmax values, and $K_{a}$ and $K_{i}$ are Michaelis-Menten constants. In steady-state

$\frac{V_{i}}{V_{a}}=r=\frac{1+ {K_{i}}/a}{1+ {K_{a}}/{(1-a)}}$ ,

which yields has solution

$a=\frac{3\times r-1-\sqrt{{9(r}^{2}-14r+9)}}{(4r-4)} .$	Eq. C4

We assume that the ratio of activities r takes the form

$r=\frac{{ACCoA}_{x}}{{CoASH}_{x}}\times\frac{{NADH}_{x}}{{NAD}_{x}}\times\frac{{ATP}_{x}}{{ADP}_{x}}\times\frac{ref\_pyr}{{PYR}_{x}}$	Eq. C5

and the PDH activity is computed

${V_{maxPDH}=a \times Vmf}_{PDH}$	Eq. C6

*Contributions from β-oxidation to NADH and acetyl-CoA:*

Acetyl-CoA is generated from two reactions in β-oxidation, Medium-chain ketoacyl-CoA thiolase

(*mkcat*) and mitochondrial trifunctional protein (*mtp*). The sum of the flux contributions are added to acetyl-CoA from the TCA cycle in Eq. C2.

$J_{\text{boxACCoA}}=\left( \frac{1}{Vx} \right)\times\frac{\left( \sum v_{mckatCn}+ \sum v_{mtpCn} \right)}{W_{x}}$	Eq. C7

NADH is generated from two reactions in β-oxidation, Medium/short-chain hydroxyacyl-CoA dehydrogenase

(*mschad*) and mitochondrial trifunctional protein (*mtp*). The sum of the flux contributions are added to NADH totals from the TCA cycle in Eq. C1.

$J_{\text{boxNADH}}=\left( \frac{1}{Vx} \right)\times\frac{\left( \sum v_{mschadCn}+ \sum v_{mtpCn} \right)}{W_{x}}$	*Eq. C8*

V_x_ is the conversion factor representing L mitochondria per mg mitochondrial protein.

Concentrations of cytoplasmic H+, Mg2+, and K+ are assumed to be fixed at buffer conditions or physiological *in vivo* values. Since the outer membrane is highly permeable to hydrogen ions and cations, we assume here

$[H^{+}]_{i}\text{ = }[H^{+}]_{c}$, $[\text{M}\text{g}^{\text{2+}}]_{i}\text{ = }[\text{M}\text{g}^{\text{2+}}]_{c}$, and $[K^{+}]_{i}\text{ = }[K^{+}]_{c}$ .

Running the Model

Complete Matlab (v.2019b, Mathworks, Natick, MA) code to reproduce figures presented here can be found at:

<https://github.com/pfizer-opensource/mitochondria-metabolism>

ReadMe.MD provides descriptions of each of the model files and instructions for usage. Briefly, add all files/directories in the github repository to the Matlab working directory/path. The Run_Final_Figures.m script will take ~7 hours to reproduce Figs. 3-6. The Cell_dxdt file contains the differential equations, the setup.m file contains initial conditions, and the PDHphosdephos.m file describes the phenomenological regulation over PDH responsible for substrate flexibility. In all cases, changes are made acutely, and the model is run to steady-state. Results are reported as a function of the free energy of ATP hydrolysis (-∆G_ATP_).
